# Maternal α-cypermethrin and permethrin exert differential effects on fetal growth, placental morphology, and fetal neurodevelopment in mice

**DOI:** 10.1101/2025.03.16.643434

**Authors:** Benjamin A. Elser, Benjamin Hing, Samuel Eliasen, Malik A. Afrifa, Naomi Meurice, Farzana Rimi, Michael Chimenti, Laura C. Schulz, Michael E. Dailey, Katherine N. Gibson-Corley, Hanna E. Stevens

**Author notes:** Corresponding author: Hanna E Stevens, MD, PhD Department of Psychiatry The University of Iowa 1330 Pappajohn Biomedical Discovery Building Iowa City, Iowa 52242.

## Abstract

Pyrethroid insecticides represent a broad class of chemicals used widely in agriculture and household applications. Human studies show mixed effects of maternal pyrethroid exposure on fetal growth and neurodevelopment. Assessment of shared pyrethroid metabolites as a biomarker for exposure obscures effects of specific chemicals within this broader class. To better characterize pyrethroid effects on fetal development, we investigated maternal exposure to permethrin, a type I pyrethroid, and α-cypermethrin, a type II pyrethroid, on fetal development in mice. Pregnant CD1 mice were exposed to permethrin (1.5, 15, or 50 mg/kg), α-cypermethrin (0.3, 3, or 10 mg/kg), or corn oil vehicle via oral gavage on gestational days (GD) 6-16. Effects on fetal growth, placental toxicity, and neurodevelopment were evaluated at GD 16. Cypermethrin, but not permethrin, significantly reduced fetal growth and altered placental layer morphology. Placental RNAseq analysis revealed downregulation of genes involved in extracellular matrix remodeling in response to α-cypermethrin. Both pyrethroids induced shifts in fetal dorsal forebrain microglia morphology from ramified to ameboid states; however, effects of α-cypermethrin were more pronounced. The α-cypermethrin transcriptome of fetal dorsal forebrain implicated altered glutamate receptor signaling, synaptogenesis, and c-AMP signaling. Coregulated gene modules in individual placenta and fetal dorsal forebrain pairs were correlated and overlapped in biological processes characterizing synapses, mitotic cell cycle, and chromatin organization, suggesting placenta-fetal brain shared mechanisms with α-cypermethrin exposure. In summary, maternal type II pyrethroid α-cypermethrin exposure but not type I pyrethroid permethrin significantly affected placental development, fetal growth, and neurodevelopment, and these effects were linked.

## Introduction

The pyrethroid class of insecticides is used in a variety of agricultural and household applications. Exposure to these chemicals is widespread, as pyrethroid metabolites are detected in the urine in 46%-79.8% of the general US population (Tang et al. 2018). In mammals, pyrethroids interact with a broad range of molecular targets, including voltage-gated sodium channels, voltage-gated calcium channels, chloride channels, and ATPases (Field et al. 2017; Kakko, Toimela, and Tahti 2003). Within this broad class, pyrethroids are classified either type I or type II depending on the presence or absence of an α-cyano group attachment to the molecule. Previous work has demonstrated that the addition of the α-cyano group leads to significant alterations in a pyrethroid’s effects on ion channels. In the context of human exposures, permethrin (a second-generation type I pyrethroid) is the most frequently used chemical in this class. However, the type II pyrethroid cypermethrin is the most frequently detected in food. The main route of exposure to these insecticides in urban areas is through the dietary intake of fruits and vegetables (Tang et al. 2018).

In comparison to previous classes of insecticides, pyrethroids are generally believed to be safe for use by the majority of the population. However, exposure to pregnant women may be of potential concern, as recent epidemiological studies have demonstrated links between exposure and adverse developmental effects (Elser, Hing, and Stevens 2022). Pyrethroid metabolites in maternal urine are negatively correlated with motor control, social adaptation, and intelligence in their children (Xue et al. 2013). Prenatal exposure to pyrethroids is also linked to increased rates of adverse behavioral outcomes and impairments in executive functioning in children (Coker et al. 2018; Watkins et al. 2016; Furlong et al. 2017). In addition, increased proximity of pyrethroid application sites to the maternal residence during pregnancy is associated with increased risk for developmental delay and autism spectrum disorders in children (Shelton et al. 2014). Several studies have also demonstrated significant correlations between maternal urinary concentrations of pyrethroid metabolites and decreased infant birth weight (Ding, Cui, et al. 2015; Hanke et al. 2003). Other studies have found that reduced fetal growth resulting from maternal pyrethroid exposure persists throughout childhood, with significant correlations between maternal urinary pyrethroid metabolites and lower BMI-for-age and weight-for-height outcomes in children (Coker et al. 2018). In the context of these findings, there is increased concern regarding reports noting increased levels of exposure in US pregnant women following restrictions on the residential use of organophosphate alternatives (Williams et al. 2008).

In contrast to the previously mentioned studies that demonstrated an increased risk for impairments in neurodevelopment and fetal growth, there have also been a considerable number of studies that have found no significant effects of prenatal pyrethroid exposure on these endpoints. Understanding potential factors that may lead to disagreement between these studies is therefore of critical importance for proper risk assessment. Due to the rapid rate of metabolism of pyrethroids, a limitation of many of these studies is that the assessment of pyrethroid exposure is often performed by the measurement of non-specific urinary metabolites such as 3-PBA and DCCA. Many of the measured metabolites are common to a wide range of chemicals within the broader pyrethroid class, which have varying effects and potencies on a range of molecular targets.

To better understand the effects of pyrethroid insecticides on fetal growth and neurodevelopment, we sought to compare the effects of two commonly used pyrethroids, permethrin and α-cypermethrin, representing the respective type I and type II sub-classifications. Both pyrethroids were investigated for their comparative effects on neurodevelopment and fetal growth following exposure throughout gestation in pregnant CD-1 mice. In the context of previous work suggesting that direct transfer of pyrethroid insecticides to fetal circulation is relatively low (Elser et al. 2022), particular emphasis was placed on the evaluation of indirect modes of action relevant to fetal growth and neurodevelopment. In this respect, given the links between low birth weight and placental insufficiency, our study included an extensive evaluation of placental toxicity for each pyrethroid. Although few studies have previously examined the effects of maternal pyrethroid insecticide exposure on placental development, effects were expected given the role that relevant molecular targets for pyrethroids play in the active transport of nutrients and regulation of growth factor/hormone synthesis. Such targets include voltage-gated calcium channels, maxi chloride channels, and plasma membrane ATPases (Vallejos and Riquelme 2007; Bernucci et al. 2006).

## Materials and Methods

### Chemicals

Analytical grade Permethrin (white solid; > 99% purity; 58.1% Trans-Isomer, 41.7% Cis-Isomer; PESTANAL, Sigma Aldrich; InChI Key: RLLPVAHGXHCWKJ-UHFFFAOYSA-N) was used for the animal studies. Neat α-cypermethrin (white solid; 98% purity; Toronto Research Chemicals, North York, Canada; InChI Key: KAATUXNTWXVJKI-NSHGMRRFSA-N) for the animal studies was purified by recrystallization in isopropanol to > 99% purity. Corn oil (vehicle) was purchased from Sigma Aldrich (Catalog No. C8267).

### Preparation of Dosing Solutions

As previously described, α-cypermethrin and permethrin dosing solutions were prepared by first dissolving each neat compound in acetone (Fisher Chemical, Certified ACS) (volume of acetone equivalent to 2% of the total volume of corn oil added), followed by evaporation of the solvent to leave a film coating the tube (Elser et al. 2022). Compounds were then dissolved in corn oil, and solutions were stored at −20 °C overnight in-between dosing. A similar process was followed for the preparation of vehicle solutions to control for the effects of residual acetone. Solutions were brought to room temperature and vortexed before dosing. Solutions were remade every 2 to 3 days to ensure minimal degradation of the compounds.

### Animals and Treatments

Experiments were conducted with approval from the Institutional Animal Care and Use Committee of the University of Iowa. CD1-IGS female mice (7-9 weeks old) were purchased from Charles River and were mated with GAD67-GFP*^+/−^* knock-in males bred on a CD1 background from our colony. GAD67-GFP*^+/−^* knock-in mice allowed for visualization of GABAergic neuronal progenitors within the fetal brain, as previous work by our lab has demonstrated a delay in tangential migration of these cells in response to short-term α-cypermethrin exposure (Elser et al. 2020). Gestational Day 0 (GD 0) was determined upon detection of a vaginal plug, and pregnant dams were singly housed from GD 0-16.5. Mice were housed in cages on a 12 h light/dark cycle with free access to food and water.

Fetuses from a total of 70 pregnant dams were evaluated following exposure to α-cypermethrin (0.3, 3, or 10 mg/kg/day), permethrin (40:60 Cis:Trans) (1.5, 15, or 50 mg/kg/day), or corn oil vehicle via oral gavage from GD 6-16 (n=10 per treatment group). Two dams exposed to 10 mg/kg/day α-cypermethrin were found dead on GD 16 prior to scheduled tissue collection; fetuses from these dams were not evaluated. Two additional replacement dams were treated with 10 mg/kg/day α-cypermethrin to allow for evaluation of a total of 10 litters within this group. Dams received a final dose of vehicle or pyrethroid 90 minutes prior to tissue collection to ensure high blood levels of the parent compounds for evaluation of maternal and fetal tissue distribution. Serum and tissue levels of cypermethrin and permethrin for all animals in this study can be found in a previously published paper (Elser et al. 2022).

The low doses for α-cypermethrin and permethrin were chosen to be just above the human acceptable daily intake (ADI) doses (0.02 mg/kg for α-cypermethrin and 0.05 mg/kg for permethrin). Human ADI doses were multiplied by a conversion factor of 12.3 to arrive at equivalent doses in mice (Nair and Jacob 2016). Within this study, these low doses correspond to average GD 16 maternal serum levels 90 minutes post-dose of 170.2 ng/mL and 256.8 ng/mL for cypermethrin and permethrin, respectively (Elser et al. 2022). Levels at this time point may reflect a degree of accumulation following repeated dosing, as previously reported (Shiba et al. 1990). Relatively higher doses of permethrin (40:60 Cis:Trans) were justified based on the ADI levels and higher level of toxicity expected for an isomer-enriched pyrethroid formulation like α-cypermethrin.

On GD 16.5, dams were anesthetized via an i.p. injection of ketamine/xylazine prior to decapitation. Maternal trunk blood was collected, allowed to clot, and then centrifuged to collect serum. The uterus was removed, rinsed with saline, weighed, and placed on ice for dissection. Maternal brain and liver were also collected, with separate samples from each tissue either fixed with 10% neutral buffered formalin or snap-frozen for chemical analysis. Amniotic fluid was collected via puncture of the amniotic sac using a 25-gauge needle and pooled across the litter. Ovaries were removed and snap-frozen for chemical analysis.

Fetuses and placentas were removed from the uterus, rinsed with saline, and then weighed with uterine position recorded. Fetuses were then quickly decapitated and GFP-positive heads were drop-fixed in 4% PFA. Placentas for RNAseq and histology (taken from GFP-positive offspring) were cut down the midline, with half placed in RNAlater after removing the decidua and the other half placed in 10% neutral buffered formalin with the decidua intact. GFP-negative brains were either snap-frozen whole for oxidative-stress measurements or submerged in RNAlater and micro-dissected for gene expression analysis of specific brain regions. Tissues submerged in RNAlater were subsequently frozen on dry ice and stored at −80°C prior to RNA extraction. GFP-negative placentas were snap-frozen whole for cytokine/growth factor ELISAs and oxidative stress assays. Following decapitation, fetal bodies were snap-frozen and utilized for analytical chemistry as detailed in a previously published paper (Elser et al. 2022). Fetal sex was determined by PCR genotyping as previously described, using primers for *Jarid1c* and *Jarid1d* (Clapcote and Roder 2005).

### RNAseq sample preparation

RNA was isolated using a RNeasy Plus Mini Kit (Qiagen) from one placenta per litter for vehicle, α-cypermethrin (0.3, 3, and 10 mg/kg), and 50 mg/kg permethrin treatment groups (n=10 per group, 5 males and 5 females per treatment from distinct litters, 50 total samples). Lower doses of permethrin (1.5 and 15 mg/kg) were not included in our placental RNAseq analysis due to an absence of effect for permethrin on fetal growth or placental layer morphology. Decidua was removed from the placentas to reduce the contribution of maternal tissue to placental gene expression that may overshadow sex-specific effects.

In addition to placental samples, RNA was isolated from the dorsal forebrain of one fetus per litter from vehicle and 3 mg/kg α-cypermethrin treatment groups (n=10 per group, 5 males and 5 females per treatment from distinct litters, 20 total samples). 3 mg/kg α-cypermethrin was chosen for comparison to control samples to avoid confounding effects of maternal toxicity observed at the 10 mg/kg α-cypermethrin dose.

Litters within treatment groups for which a male or female sample was selected were balanced by fetal body weight to control for variations in response to treatment that may influence findings regarding sex-specific effects. To control for uterine position effects, placentas from the position closest to the ovaries were not included for RNAseq analysis.

cDNA for direct quantitation of gene expression was synthesized using the AMV Reverse Transcriptase cDNA Synthesis Kit (M0277S). qPCR was performed with Power SYBR Green Master Mix (Thermo Fisher Scientific, Warrington, UK) and primers using an Applied Biosystems Model 7900HT instrument. The relative expression of each gene as compared to housekeeping gene expression was calculated using the ΔΔCt method. *Sdha* was used as a housekeeping gene for placenta samples and *Gapdh* was used as a housekeeping gene for fetal brain samples (primer sequences are listed in Supplemental Table 1).

### RNA sequencing

Transcription profiling using RNA sequencing was performed by Psomagen (Rockville, MD). Prior to library preparation, samples were quantified using fluorimetry and RNA quality was assessed using the Agilent BioAnalyzer 2100 (Agilent Technologies, Santa Clara, CA), generating an RNA Integrity Number (RIN). Useable samples were required to have a mass ≥ 1 µg, with a RIN > 8. RIN values for all RNAseq samples ranged from 8.9-9.7. Next, 500 ng of the DNase I-treated total RNA was used to enrich for polyA containing transcripts using oligo(dT) primers bound to beads. The enriched RNA pool was then fragmented, converted to cDNA and ligated to sequencing adaptors containing indexes using the Illumina TruSeq stranded mRNA sample preparation kit (Cat. #RS-122-2101, Illumina, Inc., San Diego, CA). The molar concentrations of the indexed libraries were measured using the 2100 Agilent Bioanalyzer (Agilent Technologies, Santa Clara, CA) and combined equally into pools for sequencing. The concentration of the pools was measured using the Illumina Library Quantification Kit (KAPA Biosystems, Wilmington, MA) and sequenced on the Illumina HiSeq 4000 genome sequencer using 150 bp paired-end SBS chemistry.

### Placenta RNAseq analysis

Differential expression analysis of placenta RNAseq samples was conducted by the Bioinformatics Division of the Iowa Institute of Human Genetics (IIHG, Iowa City, IA). Reads were processed with the *‘bcbio-nextgen.py’* open-source informatics pipeline developed primarily at Harvard Chan Bioinformatics (v.1.0.8) running on the Argon HPC resource at the University of Iowa. This pipeline includes ‘best practices’ approaches for read quality control, read alignment and quantitation. The ‘bcbio-nextgen.py’ pipeline was run in “RNA-seq” mode with the ‘mm10’ key as the selected genome build (internally referencing Ensembl GRCm38.p6 v94, GENCODE M19). The pipeline aligned reads to the GRCm38 genome using the splice-aware, ultra-rapid hisat2 aligner (2.1.0) and concurrently quantified reads to the GENCODE M19 transcriptome using the ‘salmon’ (0.9.1) aligner. Qualimap (2.2.2), a computational tool that examines hisat2 BAM alignment files, was used to examine the read data for quality control. Sequence quality scores passed basic checks, and sequence duplication rates were within acceptable parameters. Salmon-derived transcript quantifications (TPM) were imported and summarized to estimated counts at the gene level using *tximport* (1.12.3) in Rstudio, as described in the best-practices DESeq2 vignette (https://bioconductor.org/packages/release/bioc/vignettes/DESeq2/inst/doc/DESeq2.html).

Genes with fewer than 5 estimated counts across all samples were pre-filtered from downstream analysis, as per recommended procedure. Sex, breeder male, uterine position relative to cervix, and litter size were determined to significantly affect gene expression and were included as covariates in our analysis. Differential gene expression analysis was conducted with *DESeq2*(1.24.0) on estimated gene-level counts. An FDR of 10% and logFC of 0.5 was set as a cutoff for differential expression (DEGs). The identified differentially expressed genes (DEG)s were further analyzed to determine the gene ontologies, networks, and pathways represented. Pathway analysis of placental gene expression changes was conducted using Advaita Bioinformatics iPathwayGuide commercial software.

### Dorsal forebrain RNAseq analysis

Illumina adapter sequences were trimmed using TrimGalore (v0.5.0) with Cutadapt (v2.10). Using Kallisto (v0.46.1) (Bray et al. 2016) with options –bias –bootstrap-samples=200 –rf-stranded, data was efficiently quasi-mapped (∼92% mapping efficiency) to the mouse transcriptome, GENCODE version M25 GRCm38 (mm10), and transcript abundance was quantified. Data were analyzed in RStudio using Sleuth (Pimentel et al. 2017). Transcript abundance was normalized to gene length for gene level analysis. A principal component analysis approach was used to assess for variance between samples and Grubb’s test was used to identify outliers resulting in the removal of 1 sample. The resultant data was analyzed using a linear model where the full model consists of scaled reads per base as the response variable and treatment group and sex as the explanatory variables. Given that previous studies have shown that uterine position relative to the cervix and litter size can affect placental blood flow and nutrient availability to the fetus (Raz et al. 2012), these variables were included in the model. Similarly, as paternal genetics contribute to fetal development (Meng and Groth 2018; Tesarik 2020), breeder males used for the experiment were also included in the model. Unwanted factors were estimated using the RUVg function (k = 2) from RUVseq (Risso et al. 2014) and included in the model. Using the reduced model which excludes treatment condition, differential gene expression was assessed using likelihood ratio test as previously described (Pimentel et al. 2017) with a false discovery rate (FDR) < 0.05. Effect size was computed as log2 fold change where fold change was calculated from the mean normalized counts (i.e. scaled reads per base) of the α-cypermethrin group divided by the mean normalized counts of the control group for each gene. Genes were considered differentially expressed if they have a log2 fold change of at least 0.1 with FDR < 0.05. Pathway analysis of fetal brain gene expression changes was conducted using Ingenuity Pathway Analysis (IPA).

### Weighted gene correlation network analysis (WGCNA)

Weighted gene correlation network analysis (WGCNA) was performed using the R package WGCNA (Langfelder and Horvath 2008). One cross phenotype analysis evaluated the relationship of placental differential gene expression with other fetal phenotypes, control and 10 mg/kg α-cypermethrin data. For a second cross tissue WGCNA, placenta and fetal brain differential gene expression data from the same fetus (control and 3 mg/kg) were used for the analysis. Briefly, normalized TPM data was log-transformed using log2(x+1). Since genes with low expression and low variation can contribute to noise in the system, the function filterGenes from the R package DGCA (McKenzie et al. 2016) was used to remove lowly expressed genes (below 25^th^ percentile) and low-varying genes (below 25^th^ percentile of coefficient of variation). A correlation matrix was created from pairwise Pearson’s correlation between expression values. This matrix was then used to calculate a signed adjacency matrix by raising the correlations to a soft threshold power to achieve a scale-free topology index (R^2^) of > 0.8 according to the scale-free topology criterion (Horvath 2011). For samples used for cross phenotype analysis, a soft threshold power of 12 was used. For cross tissue analysis, a soft threshold power of 12 was used for fetal brain and 14 for placenta. Subsequently, the adjacency matrix was used to calculate a topological overlap (TOM) dissimilarity matrix, which was used to construct a dendrogram of genes. Modules of co-expressed genes were defined by the cutreeDynamic function with a minimum cluster size of 30. The function mergeCloseModules (cutHeight = 0.25) was used to merge similar modules. For each module, a module eigengene (ME) was calculated corresponding to its first principal component. Pearson’s correlation was performed between the MEs of placenta and fetal brain for cross tissue analysis. Pearson’s correlation between MEs and α-cypermethrin exposure or fetal phenotypes was also performed, and the p-values were corrected for multiple testing by false discovery rate (FDR). Hub genes defined as genes with a Gene Significance ≥ 0.2 (correlation between gene expression values across samples and trait of interest) and Module Membership ≥ 0.7 (correlation between gene expression values across samples and ME) were used for further analysis.

The hub gene list from WGCNA was loaded into the STRING database and high confidence protein interactions (0.7) was identified. Enrichment of protein-protein interaction, gene ontology and pathway analysis were performed using the analysis function from the STRING database (Szklarczyk et al. 2019). Enrichment testing of differentially expressed genes was performed using one tailed Fisher’s exact test in R.

### Differential co-expression analysis

Differential gene co-expression analysis was performed using the R package DGCA (McKenzie et al. 2016). Briefly, Pearson’s correlation was calculated using TPM values for gene pairs and compared different treatment conditions. P-values were corrected for multiple testing by false discovery rate. To evaluate the probability of observing 3 or more differentially co-expressed genes in a network of 10 genes, the number of differential co-expressed genes were calculated from a random sample of the same number of genes 1000 times. The P-value was calculated as the ratio of the number of times 3 or more genes were observed to be differentially co-expressed relative to 1000 observations.

### Placental layer morphology

Layer morphology was evaluated in two H&E-stained placental sections per litter (one male and one female when available) (n=10 per group, 140 total placentas). Areas of the labyrinth and junctional zones were outlined using a Zeiss Axiocam microscope coupled with a computer using StereoInvestigator (MBF Biosciences) and reported as percent of the total area. Layer thickness was measured at three random positions in each placental section and averaged as an alternative method for assessing layer size. Sinusoidal area measurements were performed using ImageJ. One image per placenta at 20x magnification was taken at random in the labyrinth zone for assessment of sinusoidal area. Placental sinusoidal area was determined using image thresholding in ImageJ to identify the combined areas of white space and red blood cells within the sinusoid. Sinusoidal area was reported as a percentage of total area.

Stereological cell counting within the junctional zone was performed using the unbiased fractionator approach (StereoInvestigator, MBF Biosciences) in vehicle and pyrethroid-exposed placentas. Cell counts were performed using a 200 x 150 µm counting frame on an 800 x 400 µm grid with a 20x objective lens. Placentas were analyzed from 1-2 placentas per litter (one male and one female when available, n=9-10 litters/group). Nuclei of placental glycogen cells and spongiotrophoblasts were labeled within each counting frame to determine cell counts. Collagen content within the placental junctional zone was assessed after Masson’s Trichrome staining using ImageJ. Image thresholding was used to quantify areas of blue staining (collagen) as a percentage of total junctional zone area, using two images per placenta taken randomly at 20x magnification.

### Placental macrophage IHC

Placentas utilized for histopathology were fixed in formalin, paraffin-embedded, and sectioned at a 5 µm thickness. Placental Iba1 staining was performed similarly to that in the brain, with some minor alterations. Briefly, placental sections were deparaffinized and rehydrated through a xylene/ethanol gradient. Antigen retrieval was performed by boiling the slides in 10 mM EDTA pH 6 for 10 minutes as previously described (Ghatak and Combs 2014). Slides were then washed with PBS and incubated for 1 hour in blocking solution (10% horse serum, 0.1% Tween-20, and 0.2% Triton X-100 in 0.1M PBS). After blocking, slides were incubated in the primary antibody solution overnight at 4°C (Rabbit anti-Iba-1, 1:500, WAKO), followed by incubation for 1 hour at room temp with the secondary antibody (Goat anti-rabbit 594, 1:500, Molecular Probes). Slides were cover-slipped using DAPI mounting medium (4′,6-diamidino-2-phenylindole, Vector Laboratories, #H-1200).

Placental macrophages were assessed in Iba1-stained placentas from 1-2 placentas per litter (one male and one female when available). Stereological cell counting of macrophages was performed using the unbiased fractionator approach (StereoInvestigator; MBF Biosciences). Counts were performed at 40x magnification using a 500 × 300 μm grid and 200 × 150 μm counting frames.

### Multiplex cytokine ELISA

Cytokines were assessed in maternal serum and placental lysate using a Mouse Cytokine Th17 Panel A 6-Plex kit (Biorad) (n=8-10 litters per group). Lysis buffer was prepared by dissolving protease (Roche complete Mini, EDTA-free, 1183617000) and phosphatase inhibitors (Roche PhosSTOP EASYpack, 4906845001) in T-Per Buffer (Thermo Fisher, Cat no. 78510) at a ratio of 1 tablet of each inhibitor in 10 mL Buffer. Placental lysate was prepared by homogenizing ½ placenta (40-50 mg tissue) in 500 µL of lysis buffer on ice for 20 seconds.

Homogenate was then centrifuged at 10,000 xg for 5 min at 4°C, and supernatant was aliquoted and stored at −80°C for subsequent ELISA and protein analysis. Average total protein concentration of placental lysate, as determined by Bradford assay, was 9700 ng/mL. Serum and placental lysate samples were run in duplicate as outlined in the manufacturer’s guide. Samples were diluted in Bioplex sample diluent at a ratio of 1:4 for serum and 1:10 for placental lysate. The manufacturer’s protocol was followed for all samples run in duplicate.

### Fetal brain cortical volume measurements

Cortical volume measurements were performed using two fetal brains per litter (one male and one female brain when available) (n=10 per group, 140 brains total). Cortical volume was determined by measuring cortical area in 5-6 serial sections per brain (25 µm thick sections, each 20 sections apart).

### Fetal brain microglia assessment

GAD67-GFP+ fetal brains were fixed in 4% paraformaldehyde and stained for Iba1 as previously described (Elser et al. 2020). Stereological cell counting of Iba1+ microglia was performed using the unbiased optical fractionator approach (StereoInvestigator; MBF Biosciences). Counts were performed at 40x magnification using a 500 × 300 μm grid and 200 × 150 × 15 μm counting frames. Cell density was assessed stereologically using 3-4 serial coronal sections (every 20 sections apart) of the fetal cortical plate as previously described (Gumusoglu et al. 2017; Bittle and Stevens 2018). Several brains were stained with the microglial marker anti-p2ry12 (AnaSpec polyclonal rabbit anti-mouse p2ry12) using the same IHC processes to confirm the identity of Iba1+ ameboid cells as microglia.

### MDA measurement

Lipid peroxidation marker Malondialdehyde (MDA) was quantified in offspring placenta homogenate using the thiobarbituric acid reactive substance (TBARS) assay and was normalized to total protein as determined by BCA assay (Janero 1990). Briefly, one male and one female placenta from each litter (n=9-10 per group) were homogenized separately in RIPA buffer, followed by centrifugation (3000xg, 10 min, 4°C). MDA standards (0-50μM) were prepared using 1,1,3,3-tetramethoxypropane. Sample supernatant or standard and acid solution (15%TCA, 0.37%TBA, 0.25M HCl) were incubated (95°C, 45 min) and centrifuged (14,000 rpm, 5 min, 4°C). Supernatants were vortexed with n-butanol and saturated NaCl (30s), followed by centrifugation 14,000 rpm, 2 min, 4°C). Supernatant absorbance was measured in duplicate at 535 nm and 572 nm (correcting for blank levels). Corrected absorbance values (Abs 535-Abs 572) were used to calculate concentrations of MDA in each sample using the standard curve.

### Statistical analysis

The litter was treated as the statistical unit for all measurements, using one male and one female per litter when available. On average, tissues from over half of all embryos from each litter were used for analysis, dispersed across multiple experiments. Sex differences were first evaluated for main effect and interaction and if not found, male and female samples within litters were averaged. Data were analyzed by ANOVA for main effects of α-cypermethrin and permethrin separately, followed by Dunnett’s multiple comparisons test to control. Outliers identified using Grubbs’ Test were removed prior to statistical analysis. All analyses were performed with GraphPad Prism (San Diego, CA). Venn diagrams were created using InteractiVenn (Heberle et al. 2015).

## Results

### Maternal toxicity

Maternal toxicity was assessed through evaluation of maternal body weight gain, post-dose observations, liver weight and histology, and measurement of serum cytokines. Throughout the treatment period (GD 6-16), maternal weight gain (absolute or adjusted for gravid uterine weight) was unaffected by α-cypermethrin (Cyp). However, there was a statistically significant, but non-dose-responsive, increase in maternal weight gain adjusted for gravid uterine weight in the 1.5 and 50 mg/kg permethrin (Per) groups (Table 1, permethrin main effect, F=5.568, p=0.0031; Per 1.5 mg/kg post hoc, p=0.0138; Per 50 mg/kg post hoc, p=0.0277). Despite a lack of weight gain change, 2/12 mice treated with 10 mg/kg α-cypermethrin were found dead on the morning of GD 16. In addition, 3/10 surviving mice treated with 10 mg/kg α-cypermethrin exhibited neurobehavioral alterations consistent with pyrethroid-induced choreoathetosis (involuntary movements) approximately one hour after dose administration on GD 16. Mortality and neurobehavioral alterations in response to 10 mg/kg α-cypermethrin were unexpected based on previous studies in pregnant mice (Elser et al. 2020). Given that these findings were only observed in mice within the high-dose cypermethrin group and were not observed in any other dose group or controls, these effects were considered likely related to 10 mg/kg α-cypermethrin exposure. However, the possibility of mortality being secondary to inadvertent gavage error could not be completely ruled out. In comparison, there was no mortality or neurobehavioral alterations noted in any of the permethrin-treated dams.

**Table 1.** Maternal and developmental toxicity endpoints in CD1 mice exposed to α-cypermethrin or permethrin from GD 6-16.

| | α-Cypermethrin (mg/kg/day, po) | | | | Permethrin (mg/kg/day, po) | | |
| --- | --- | --- | --- | --- | --- | --- | --- |
|  | Vehicle | 0.3 | 3 | 10 | 1.5 | 15 | 50 |
| **Maternal body weight (g)** |  |  |  |  |  |  |  |
| GD 0 | 27.1 ± 0.7 | 27.5 ± 0.6 | 27.9 ± 0.6 | 26.9 ± 0.5 | 28.7 ± 0.6 | 28.5 ± 0.6 | 27.8 ± 0.8 |
| GD 6 | 29.9 ± 0.8 | 30.8 ± 0.8 | 31.2 ± 0.7 | 30.4 ± 0.5 | 32.6 ± 0.8 | 32.0 ± 0.8 | 31.8 ± 0.9 |
| GD 16 | 51.4 ± 1.3 | 51.1 ± 1.4 | 53.0 ± 1.7 | 49.6 ± 1.3 | 57.3 ± 1.8 | 52.7 ± 2.3 | 55.8 ± 1.9 |
| **Body weight gain (g)** |  |  |  |  |  |  |  |
| Treatment (GD 6-16) | 21.5 ± 1.0 | 20.3 ± 0.8 | 21.8 ± 1.1 | 19.2 ± 1.0 | 24.7 ± 1.2 | 20.7 ± 1.6 | 24.1 ± 1.1 |
| Gestation (GD 0-16) | 24.3 ± 1.0 | 23.6 ± 0.9 | 25.1 ± 1.4 | 22.8 ± 1.0 | 28.6 ± 1.3 | 24.2 ± 1.8 | 28.0 ± 1.3 |
| Corrected | 7.5 ± 0.5 | 7.9 ± 0.4 | 8.3 ± 0.7 | 7.7 ± 0.7 | 10.5 ± 0.7* | 7.4 ± 0.9 | 10.2 ± 0.7* |
| **Gravid uterine weight (g)** | 16.8 ± 0.7 | 15.6 ± 0.6 | 16.7 ± 0.8 | 15.1 ± 0.5 | 18.2 ± 0.9 | 15.8 ± 0.9 | 17.8 ± 0.9 |
| **Maternal liver weight** |  |  |  |  |  |  |  |
| Absolute (g) | 2.40 ± 0.08 | 2.38 ± 0.07 | 2.50 ± 0.07 | 2.28 ± 0.06 | 2.66 ± 0.09 | 2.60 ± 0.13 | 2.70 ± 0.10 |
| Relative (% body weight) | 4.7 ± 0.1 | 4.7 ± 0.08 | 4.7 ± 0.09 | 4.6 ± 0.09 | 4.6 ± 0.07 | 4.92 ± 0.07 | 4.83 ± 0.08 |
| **Post-implantation loss (%)** | 5 ± 2 | 5 ± 2 | 4 ± 2 | 4 ± 2 | 1 ± 1* | 2 ± 1 | 2 ± 1 |
| **Live litter size (n)** | 13 ± 1 | 13 ± 1 | 14 ± 1 | 13 ± 1 | 14 ± 1 | 13 ± 1 | 14 ± 1 |
| **Fetal body weight (g)** | 0.80 ± 0.01 | 0.76 ± 0.01 | 0.75 ± 0.02 | 0.72 ± 0.01** | 0.77 ± 0.02 | 0.79 ± 0.01 | 0.80 ± 0.01 |
| **Placental weight (mg)** | 90 ± 3 | 90 ± 2 | 88 ± 3 | 88 ± 2 | 97 ± 2 | 95 ± 2 | 97 ± 4 |

Values reported as mean ± SEM.
\*p<0.05, \*\*p<0.001; pairwise comparison to control

Given the involvement of maternal hepatic function in the regulation of fetal growth, maternal liver toxicity was evaluated in response to pyrethroid treatment. Maternal liver weight (absolute and relative to body weight) was unchanged by either treatment. Maternal liver histology from dams exposed to vehicle and high doses of α-cypermethrin and permethrin (n=5 per treatment group) was evaluated by a board-certified veterinary pathologist (KGC). No significant differences in maternal liver pathology were found between vehicle and pyrethroid-treated livers (Supplemental Figure 1).

Previous studies have implicated maternal inflammation as a causative factor in fetal growth restriction and adverse neurodevelopmental outcomes of environmental toxicants (Sauder et al. 2019; Al-Azemi, Raghupathy, and Azizieh 2017). As such, maternal inflammation was evaluated through measurement of a panel of pro- and anti-inflammatory cytokines in maternal serum (n=8-10 samples per treatment group). Interestingly, 10 mg/kg α-cypermethrin reduced serum levels of both the pro-inflammatory cytokine IL-1β by 27.7% (Cyp main effect, F=4.064, p=0.0141; post hoc p=0.0367) and the anti-inflammatory cytokine IL-10 by 34.1% relative to control (Cyp main effect, F=4.602, p=0.0081; post hoc, p=0.0177) (Table 2). Although the changes in serum cytokines were of relatively low magnitude, this finding is consistent with previous work demonstrating suppression of serum cytokines of cypermethrin-exposed pesticide applicators (Costa et al. 2013). Compared to α-cypermethrin, dams treated with 1.5 mg/kg permethrin demonstrated a 33.5% increase in serum IL-17a relative to control; however, no significant effect on IL-17a was observed at higher doses of permethrin and there were no additional effects on other cytokines in permethrin-treated dams (Table 2, Per main effect, F=3.243, p=0.0357; post hoc, p=0.0365). Given the lack of dose responsiveness, the relationship to treatment is unclear with regard to the increase in IL-17a. However, increased circulating levels of pro-inflammatory cytokines have been previously reported following long-term treatment in rats with low doses of permethrin (2.5 and 5 mg/kg) (Feriani et al. 2021).

**Table 2.** Maternal serum cytokine levels in CD1 mice exposed to α-cypermethrin or permethrin from GD 6-16 (ng/mL)

|  | **α-Cypermethrin (mg/kg/day, po)** |  |  |  | **Permethrin (mg/kg/day, po)** |  |  |
| --- | --- | --- | --- | --- | --- | --- | --- |
|  | Vehicle | 0.3 | 3 | 10 | 1.5 | 15 | 50 |
| **IL-6** | 18.1 ± 1.5 | 18.6 ± 1.4 | 20.0 ± 1.7 | 15.7 ± 1.8 | 20.6 ± 1.4 | 16.5 ± 1.4 | 22.6 ± 2.6 |
| **IFN-g** | 294.2 ± 16.4 | 325.1 ± 25.6 | 326.7 ± 23.4 | 232.0 ± 33.1 | 387.9 ± 35.8 | 276.4 ± 21.9 | 417.9 ± 68.8 |
| **IL-17a** | 1395 ± 59 | 1415 ± 112 | 1588 ± 87 | 1169 ± 107 | 1863 ± 132* | 1471 ± 105 | 1774 ± 176 |
| **IL-10** | 373.6 ± 25.8 | 381.3 ± 34.2 | 375.4 ± 30.2 | 246.1 ± 31.3* | 390.6 ± 39.7 | 362.9 ± 30.2 | 448.1 ± 51.5 |
| **IL-1B** | 26.10 ± 1.89 | 26.49 ± 1.92 | 27.20 ± 1.79 | 18.86 ± 2.17* | 27.65 ± 2.24 | 22.28 ± 1.63 | 31.05 ± 3.68 |
| **TNF-a** | 1262 ± 67.32 | 1372 ± 108 | 1460 ± 125 | 977 ± 134 | 1741 ± 190 | 1263 ± 145 | 1737 ± 241 |

Values reported as mean ± SEM.
\*p<0.05; pairwise comparison to control

### Embryo/fetal growth and viability

No effect on embryo/fetal viability was observed for either pyrethroid tested; the rate of post-implantation loss and number of live fetuses per litter were comparable to control for all treatment groups (Table 1). However, α-cypermethrin induced a dose-dependent reduction in fetal body weight. Mean fetal weights were reduced by 4.9%, 6.1%, and 10.0% relative to control for litters treated with 0.3, 3, and 10 mg/kg α-cypermethrin, respectively (Table 1 & Figure 1A, Cyp main effect, F= 4.340, p=0.0104; 10 mg/kg post hoc, p=0.0029). In comparison, no significant effect of permethrin on fetal body weight was observed.

**Fig. 1.**
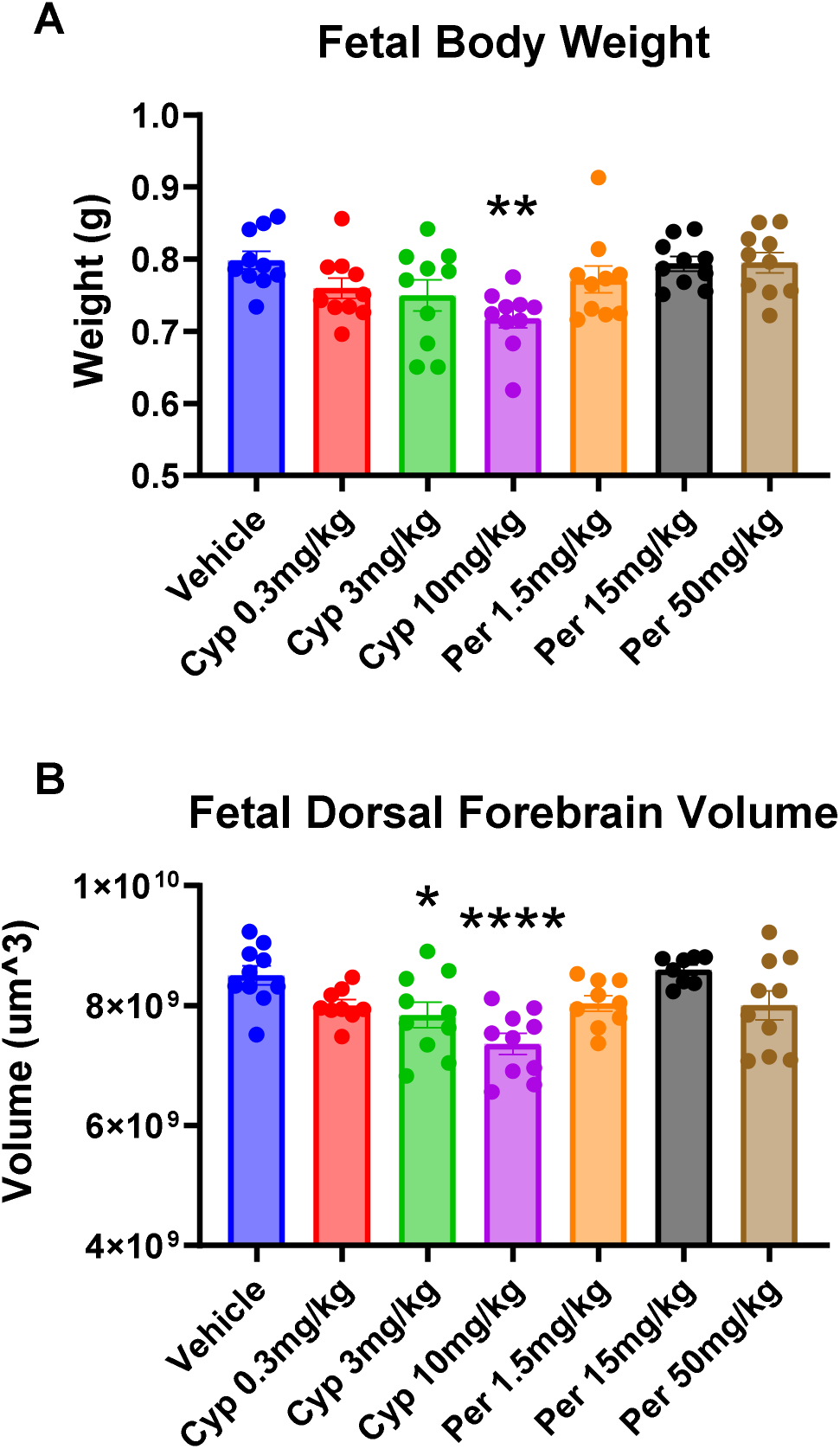
Cypermethrin induces symmetrical growth restriction in CD-1 mouse fetuses. (A) Cypermethrin, but not permethrin, reduces fetal weight. (B) Cypermethrin reduces GD 16 fetal dorsal forebrain volume. Means ± SEMs are shown. n=9-10 litters per treatment group. (*p<0.05, **p<0.01, ****p<0.0001 by post hoc tests). Cyp= Cypermethrin, Per= Permethrin.

Given the reduction in fetal weight, stereological measurements of dorsal forebrain volume were performed to determine whether brain growth was affected by treatment. Cypermethrin, but not permethrin, significantly reduced dorsal forebrain volume in a dose-dependent manner, with statistically significant effects observed at 3 and 10 mg/kg (Figure 1B, Main Effect, F=8.066, p=0.0003; 3 mg/kg Cyp post hoc p=0.0213, 10 mg/kg Cyp post hoc p<0.0001). Dorsal forebrain volume was significantly correlated to fetal body weight, suggesting that this effect is likely a result of symmetrical growth restriction, which can be the result of placental dysfunction (Supplemental Figure 3A; R=0.66, p<0.0001) (Sun et al. 2020).

### Placental Layer Morphology

No significant alterations in placental weight were observed for any treatment group (Table 1). However, placenta layer morphology was significantly altered by both 3 mg/kg (p=0.0378) and 10 mg/kg α-cypermethrin (p=0.0020), resulting in a smaller relative labyrinth zone area and a larger relative junctional zone area (Figure 2A & B, Cyp main effect, F= 4.833, p=0.0063). Cypermethrin-induced alterations in placental morphology were further confirmed by measurement of layer thickness. Mean labyrinth zone thickness was 5.6%, 10.8%, and 15.7% less than control for litters exposed to 0.3, 3, and 10 mg/kg α-cypermethrin, respectively (Supplemental Figure 2, Cyp main effect, F= 4.564, p=0.0083; Cyp 10 mg/kg post hoc p=0.0035). In addition, placentas from litters exposed to 10 mg/kg α-cypermethrin demonstrated a significant increase in junctional zone thickness (Supplemental Figure 2B, Cyp main effect, F= 3.34, p=0.0298; post hoc p=0.0140). Previous work has demonstrated that impairments in labyrinth zone development may result in fetal growth restriction by reducing blood flow to the fetus (Woods, Perez-Garcia, and Hemberger 2018). Consistent with a link between altered labyrinth zone development and fetal growth, a significant positive correlation was observed between labyrinth zone thickness and fetal body weight (Supplemental Figure 3B, r=0.56, R^2^=0.31, p<0.0001). In contrast, there was no correlation between junctional zone thickness and fetal body weight. No effects on placental layer size were observed for permethrin at any dose investigated.

**Fig. 2.**
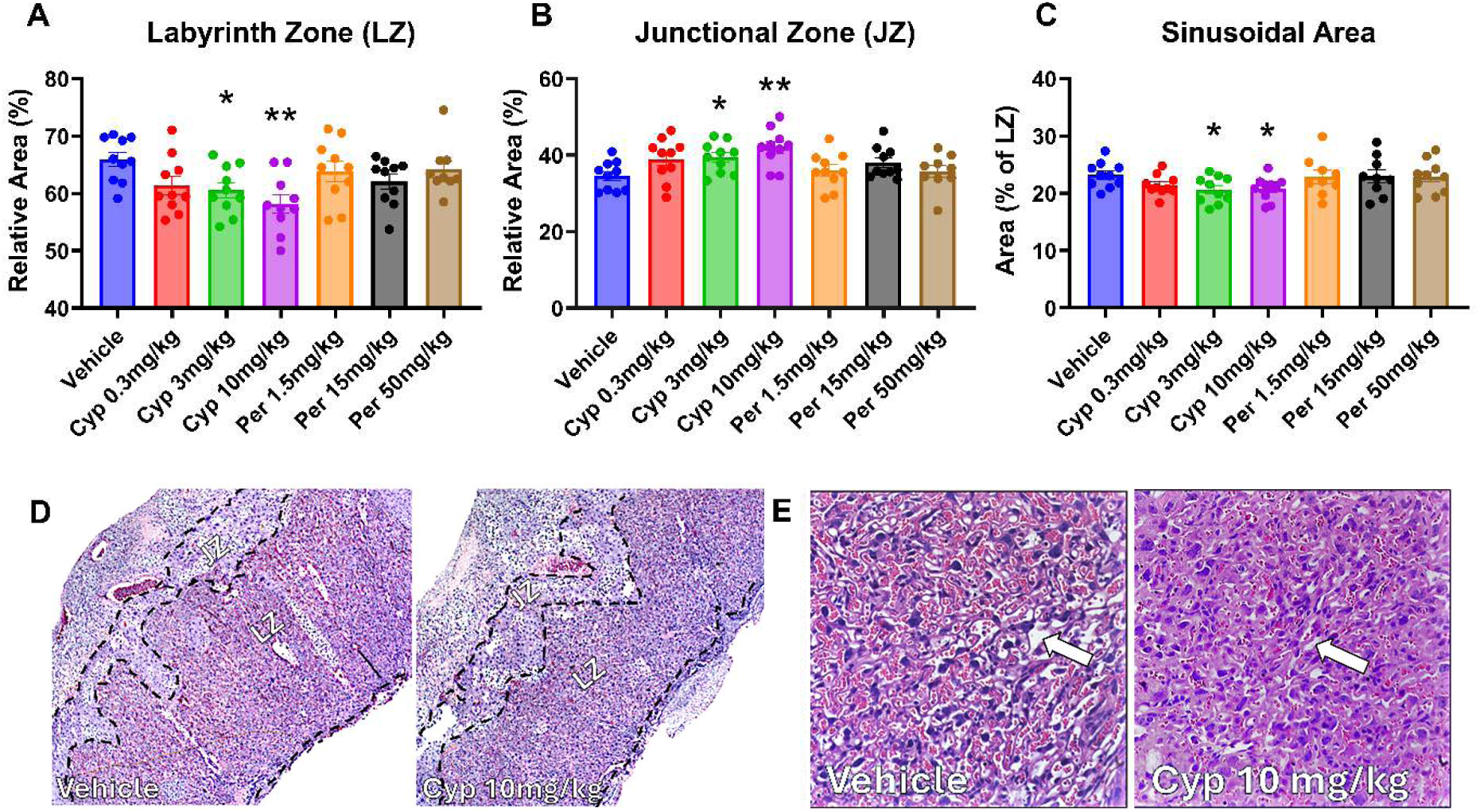
**Cypermethrin alters placental layer morphology in the GD 16 mouse placenta.**Placental labyrinth (A) and junctional zone areas (B) reported as percent of total placental area. (C) Cypermethrin reduced sinusoidal area in the labyrinth zone. (D) Representative pictures of placentas from litters treated with vehicle or 10 mg/kg cypermethrin highlighting the labyrinth and junctional zones. (E) Representative picture of the placental labyrinth zone from litters treated with vehicle or 10 mg/kg cypermethrin highlighting the difference in sinusoidal space (arrows=example sinusoids). Means ± SEMs are shown. n=9-10 litters per treatment group. (*p<0.05, **p<0.01 by post hoc tests) Cyp= Cypermethrin, Per= Permethrin.

Cypermethrin-induced alterations in placental labyrinth zone morphology were further investigated through measurement of sinusoidal area. Within the labyrinth zone, sinusoidal area was reduced by 7.7%, 11.0%, and 10.2% relative to control for litters treated with 0.3, 3, and 10 mg/kg α-cypermethrin, respectively (Figure 2C, Cyp main effect, F= 3.117, p=0.0380; Cyp 3 mg/kg post doc p= 0.0300; Cyp 10 mg/kg post hoc p=0.0340). No effect on sinusoidal area was observed for permethrin.

Given that the placental junctional zone is comprised of multiple cell types with differing functions, cell counts were performed to further characterize α-cypermethrin-induced alterations in the junctional zone. Within the junctional zone, we observed a 50.4%, 41.7%, and 74.1% increase in the number of glycogen cells relative to control for litters exposed to 0.3, 3, and 10 mg/kg α-cypermethrin, respectively (Figure 3A, Cyp main effect, F= 3.378, p=0.0204; Cyp 10 mg/kg post hoc p=0.0096). However, no significant difference in spongiotrophoblast number was observed for any of the treatment groups (Figure 3B); these findings suggested that junctional zone enlargement in response to α-cypermethrin exposure may result from a selective expansion of the glycogen cell population. No significant effects on junctional zone cell populations were observed in response to permethrin exposure.

**Fig. 3.**
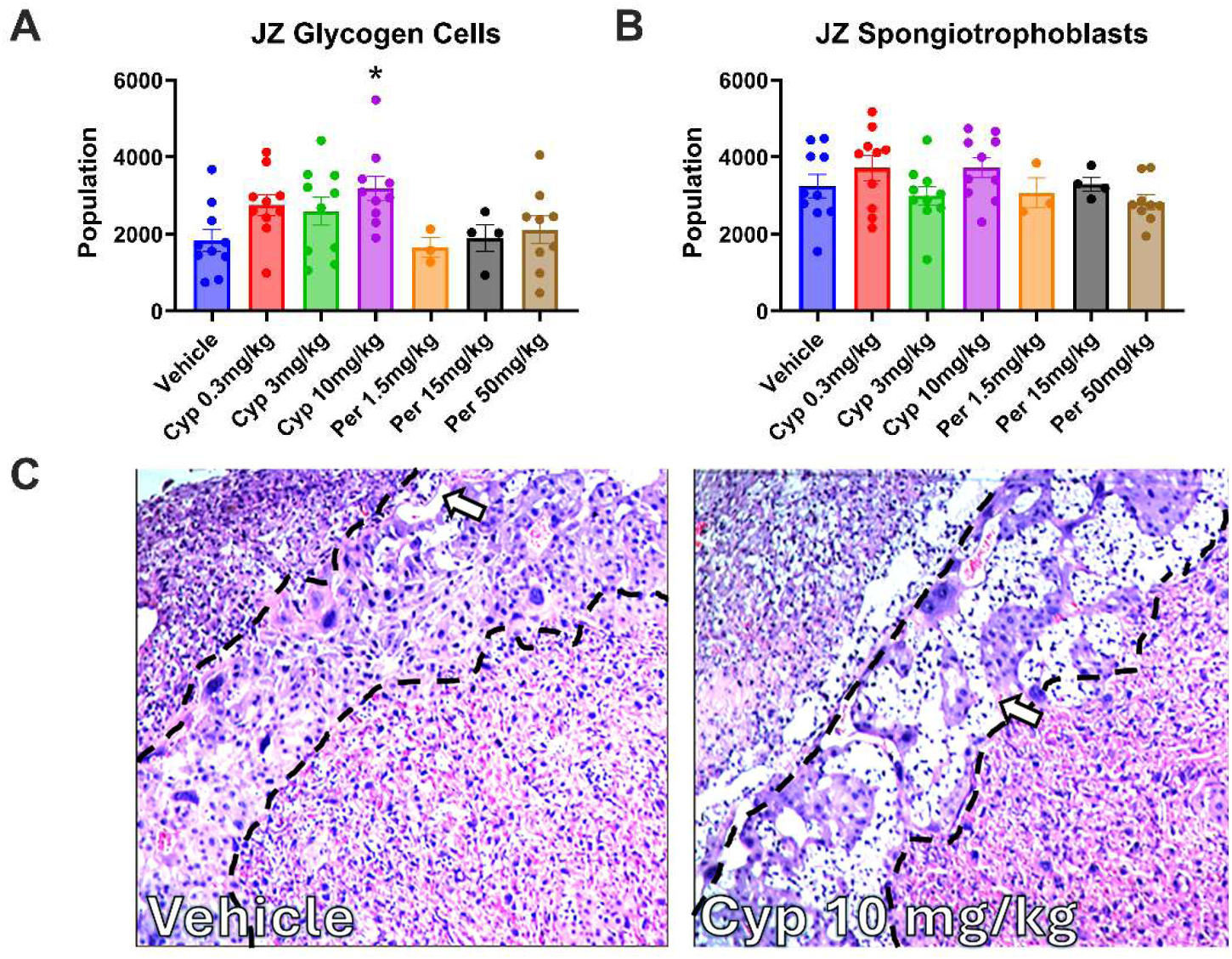
Cypermethrin increases the numbers of junctional zone (JZ) glycogen cells in the GD 16 mouse placenta. (A) Cypermethrin increased the junctional zone glycogen cell population. (B) Junctional zone spongiotrophoblast population was unaffected by either pyrethroid (C) Representative pictures demonstrating increased glycogen cell population (H&E Stain, arrows=examples of glycogen cells) in the placental junctional zone in response to 10 mg/kg cypermethrin. Means ± SEMs are shown. n=8-10 litters per treatment group. (*p<0.05 by post hoc tests). Cyp= Cypermethrin, Per= Permethrin.

### Placenta RNAseq analysis

Given the alternations in fetal growth and placental morphology noted in response to α-cypermethrin treatment, RNAseq analysis of placental samples was performed to assess the potential mechanisms by which each pyrethroid may impact placental development. Due to the lack of effects that permethrin exhibited on fetal growth or placental morphology, only 50 mg/kg permethrin was examined via RNAseq. Consistent with impacts to placental morphology, α-cypermethrin significantly altered placental gene expression across all doses tested (Figure 4A). In contrast to high dose (10 mg/kg) α-cypermethrin, high dose 50 mg/kg permethrin exhibited less changes on placental gene expression (31 differentially expressed genes (DEGs); Supplemental Fig 4). Interestingly, while placentas from dams administered 10 mg/kg α-cypermethrin (100 DEGs) demonstrated a higher number of DEGs than the lower doses, we observed a relatively higher number of DEGs at the 0.3 mg/kg dose (59 DEGs) compared to the 3 mg/kg dose (27 DEGs) (Figure 4A). In addition, male placentas (140 DEGs) showed a higher number of DEGs than females (e.g. 44 DEGs in response to 10 mg/kg α-cypermethrin (Supplemental Figure 4)). In contrast to these findings, there were no clear sex-specific differences with regard to placental layer morphology or fetal weight.

**Fig. 4.**
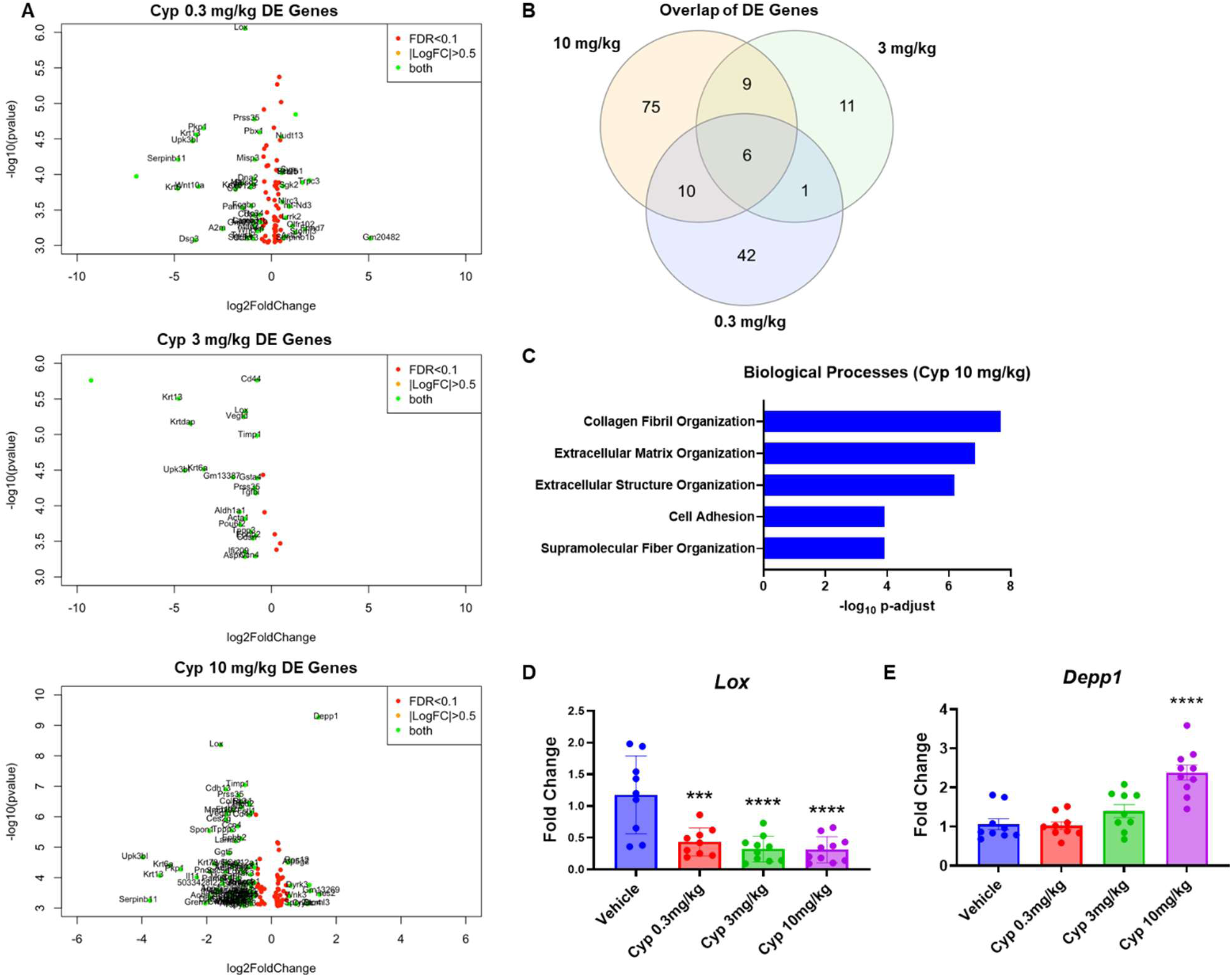
**Cypermethrin alters placental gene expression pathways involved in organization of collagen fibrils and the extracellular matrix.**(A) Volcano plots of differentially expressed (DE) genes in response to 0.3, 3, or 10 mg/kg cypermethrin. (B) Overlap of DE genes across 0.3, 3, and 10 mg/kg cypermethrin (InteractiVenn). (C) Cypermethrin altered genes involved in organization of collagen fibrils and the extracellular matrix (iPathway Analysis). (D) Cypermethrin decreased placental *Lox* gene expression via qPCR. (E) Cypermethrin increased placental *Depp1* gene expression via qPCR. Means ± SEMs are shown. n=9-10 litters per treatment group. (***p<0.001, ****p<0.0001 by post hoc tests). Cyp= Cypermethrin, Per= Permethrin.

Gene ontology analysis via iPathway demonstrated significant alterations in genes involved in the organization of the extracellular matrix and collagen fibrils in response to α-cypermethrin exposure (Figure 4B). In contrast to α-cypermethrin, gene ontology analysis of permethrin-exposed placentas revealed few statistically significant alterations in biological processes, consistent with a lack of effect on placental morphology (Supplemental Figure 5). Analysis of DEGs identified *Lox* as one of the most significantly altered genes in response to α-cypermethrin. qPCR confirmed a significant downregulation of placental *Lox* expression across all three doses of α-cypermethrin (56.2%, 67.8%, and 68.8% reduction relative to control for 0.3, 3, and 10 mg/kg α-cypermethrin, respectively) (Figure 4D, Cyp main effect, F=12.92, p<0.0001; 0.3 mg/kg post hoc p=0.0002, 3 mg/kg post hoc p<0.0001, 10 mg/kg post hoc p<0.0001). Lysyl oxidase (Lox) is an enzyme responsible for catalyzing the cross-linking of collagen and elastin in the extracellular matrix, with critical placental roles in vascular development and decidualization (Li et al. 2017; Hein et al. 2001; Xu et al. 2019; Wang et al. 2023). In addition, analysis of DEGs identified a dose-dependent increase in expression of the autophagy regulator *Depp1* in response to α-cypermethrin, which was also confirmed via qPCR (Figure 4E, Cyp main effect, F=17.46, p<0.0001; 10 mg/kg post hoc p<0.0001).

### Placental Collagen Content

Based on significant alterations in genes involved in collagen fibril organization and previous reports of increased trophoblast secretion of collagen with downregulated *Lox* expression (Xu et al. 2019), we next investigated whether placental collagen content was altered by α-cypermethrin exposure. Consistent with previous reports, placentas exposed to 10 mg/kg α-cypermethrin demonstrated a significant increase in collagen staining within the junctional zone (Figure 5A; Cyp main effect, F=4.613, p<0.01; 10 mg/kg post hoc p<0.01).

**Fig. 5.**
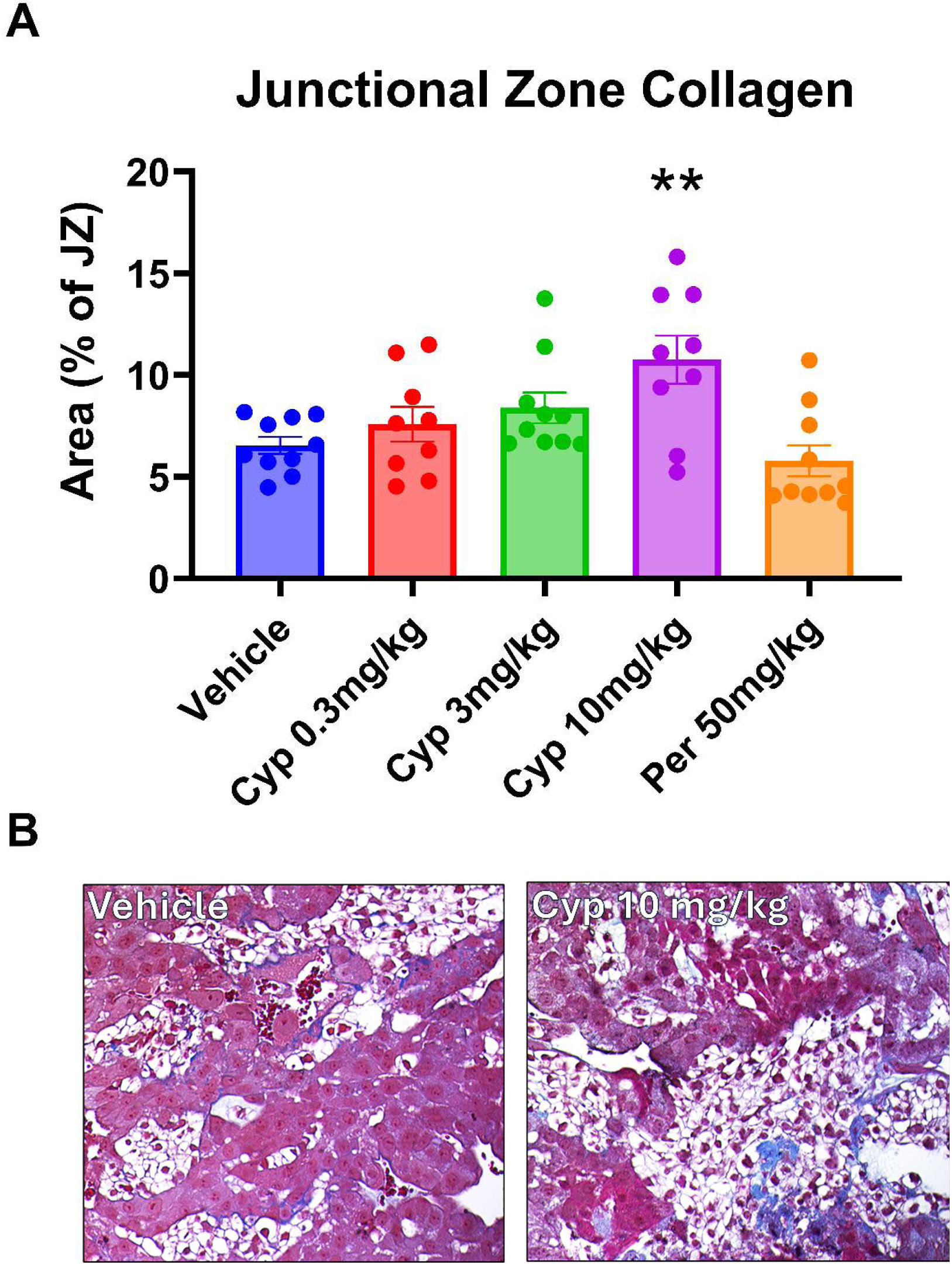
Cypermethrin increases placental collagen staining in the junctional zone. (A) Cypermethrin increased collagen content in the junctional zone as assessed by Masson’s Trichrome Stain. (B) Representative pictures of placentas stained with Massons Trichrome. Collagen is stained blue. Means ± SEMs are shown. n=8-10 litters per treatment group. (**p<0.01 by post hoc test). Cyp= Cypermethrin, Per= Permethrin.

These findings were also consistent with previous work demonstrating increased collagen deposition in cardiac tissue of rats following adult long-term treatment with either α-cypermethrin, permethrin, or deltamethrin (Feriani et al. 2021; Feriani et al. 2020; Ghazouani et al. 2020).

### Assessment of placental oxidative stress and inflammation

Upregulation of *Depp1* has been shown to occur in response to a variety of stimuli, including hypoxia/oxidative stress, fasting, and progesterone (Chen et al. 2011). Given that previous reports have suggested a link between impaired labyrinth zone development and hypoxia/oxidative stress, we investigated whether α-cypermethrin increased markers of oxidative stress in placental tissue (Eskild, Strøm-Roum, and Haavaldsen 2016). In contrast to our previous work demonstrating an increase in placental malondialdehyde content following short-term exposure to α-cypermethrin and restraint stress, neither pyrethroid induced a significant alteration in placental malondialdehyde within this study (Supplemental Figure 6A). Given that oxidative stress may be compensated for through upregulation of antioxidant enzyme activity, thioredoxin reductase activity was assessed in vehicle-treated and α-cypermethrin-exposed placentas. An increase in thioredoxin reductase activity was noted in response to 3 mg/kg cypermethrin, but the lack of dose dependence suggests interpreting this difference with caution (Supplemental Figure 6B). Altogether, our results do not suggest that α-cypermethrin exposure induced a significant degree of oxidative stress in GD 16 placental tissue.

Assessment of inflammatory indicators in placenta also showed little alteration (Supplemental Figure 7). Overall, decreased placental Iba1+ macrophage density was observed in male placentas in response to 0.3 mg/kg α-cypermethrin; however, no significant effect was observed at higher doses in males or any dose in females. Placental macrophage density was also unchanged in response to 50 mg/kg permethrin (Supplemental Figure 7A). In addition, no notable effect of α-cypermethrin or permethrin was observed in regard to placental cytokine production. Although an increase in placental IL-1β was observed in the 0.3 mg/kg α-cypermethrin group, no effect was observed at higher doses (Supplemental Figure 7B, Cyp main effect, F=4.096, p=0.0068; post hoc p=0.019).

### Fetal brain microglia

Microglia, inflammatory cells in the brain, have been strongly implicated in the pathogenesis of neurodevelopmental disorders and play a significant role in neural progenitor proliferation, synaptogenesis, myelination, and synaptic pruning (Bordeleau et al. 2019). Alterations in microglial activation state have also been implicated in models of growth restriction (Miller, Huppi, and Mallard 2016). In addition, previous work by our lab demonstrated that a three day exposure to α-cypermethrin and restraint stress alters microglial morphology in the GD 14 mouse brain (Elser et al. 2020). As such, the present study investigated fetal microglia development following pyrethroid exposure GD 6-16.

Consistent with previous findings, α-cypermethrin exposure induced a dose-dependent shift in dorsal forebrain microglia from a more developed ramified morphology towards a less-developed ameboid morphology (Figure 6C, Cyp main effect, F=9.369, p=0.0001; 0.3 mg/kg Cyp post hoc p=0.0170, 3 mg/kg Cyp post hoc p=0.0015, 10 mg/kg Cyp post hoc p<0.0001). Two-way ANOVA revealed a significant interaction between sex and α-cypermethrin exposure for the percentage of ramified microglia. When data were split by sex, all doses of α-cypermethrin in males demonstrated significant decreases in the percentage of ramified microglia, whereas females were only significantly affected at 3 and 10 mg/kg (Supplemental Figure 8). In comparison, permethrin exposure resulted in a much smaller shift towards the ameboid state, with only the 50 mg/kg permethrin dose demonstrating a significant increase in this morphology relative to control (Figure 6C, Per main effect, F=3.293, p=0.0321; 50 mg/kg Per post hoc p=0.0159). Lastly, both 3 mg/kg (p=0.0165) and 10 mg/kg α-cypermethrin (p=0.0021) led to significant increases in the phagocytic multivacuolated state of microglia (Figure 6C, Cyp main effect, F=7.905, p=0.0004). No significant effect of permethrin was found for multivacuolated microglia (Figure 6C).

**Fig. 6.**
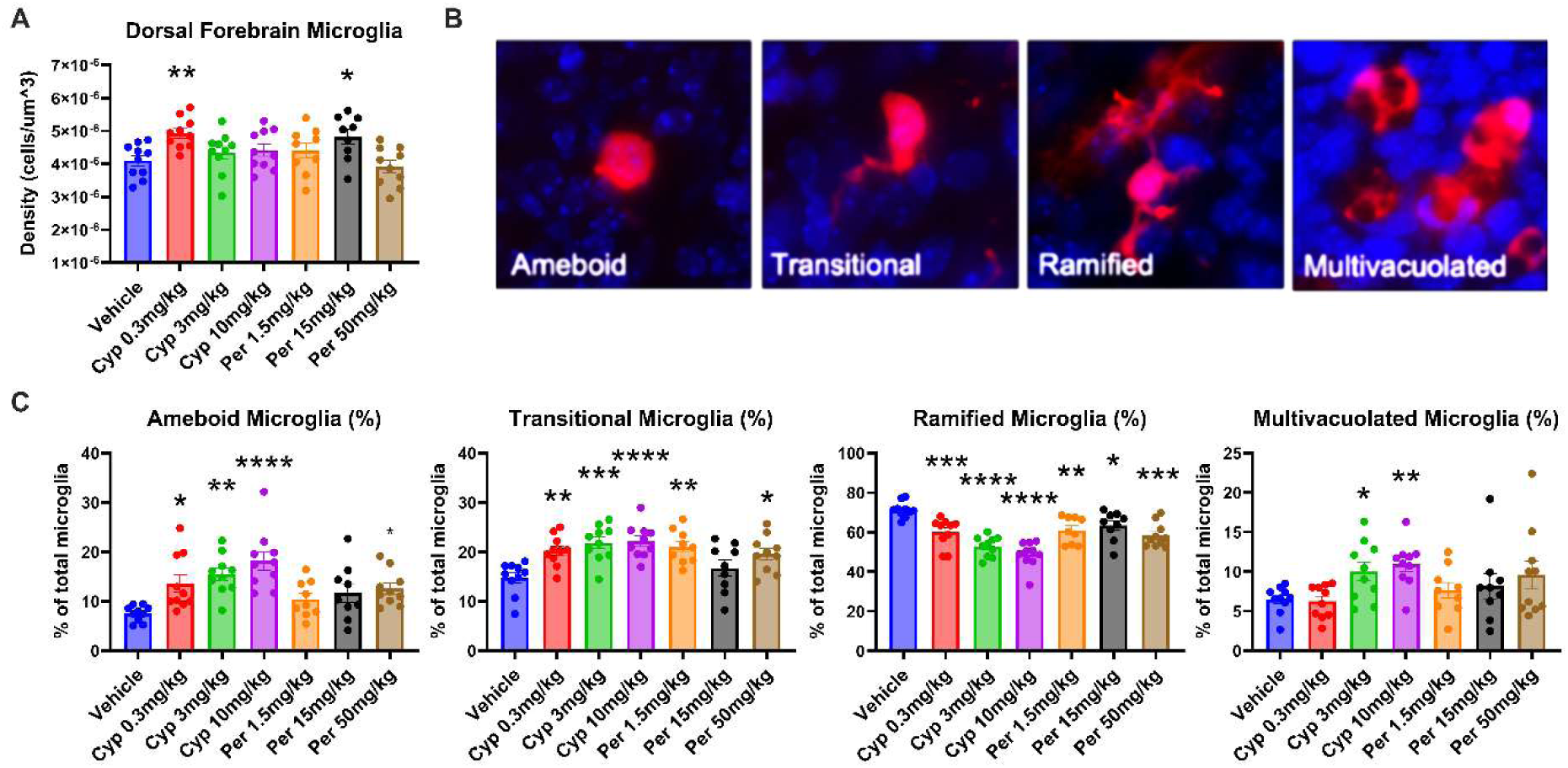
Alterations in fetal brain microglia development following gestational exposure to pyrethroids. (A) Total microglia density in the GD16 fetal brain. (B) Representative pictures of GD 16 microglia morphologies immunostained with Iba1 in red; nuclear DAPI is in blue: 1) Ameboid, with no processes and normal nucleus, 2) Transitional, with one process and normal nucleus, 3) Ramified, with two or more processes and normal nuclei, 4) Multivacuolated, with multiple vacuoles and/or pyknotic nuclei (high magnification). Means ± SEMs are shown. n=9-10 litters per treatment group. (*#*p<0.01, *p<0.05, **p<0.01, ***p<0.001, ****p<0.0001 by post hoc tests). Cyp= Cypermethrin, Per= Permethrin.

Iba-1 positive ameboid cells may reflect infiltrating macrophages rather than microglia. However, similar total microglia density across groups, without an increased amount due to infiltrating cells, (Fig. 7A), supports that ameboid cells were microglia. Additionally, ameboid cells were similarly labeled with P2RY12 between the groups, supporting their identity as microglia rather than infiltrating macrophages (Supplemental Figure 9). This shift towards ameboid morphology may represent either a developmental delay or an activated state of these cells. However, expression of microglial activation genes (Walker and Lue 2015) (*Cd68*, *Apoe*, and *Il-1b*) or the resting state gene (*P2ry12)* were unchanged between vehicle and 10mg/kg α-cypermethrin treated brains (Supplemental Figure 9), suggesting that this shift in microglia morphology may reflect a developmental delay rather than an activated state.

**Fig. 7.**
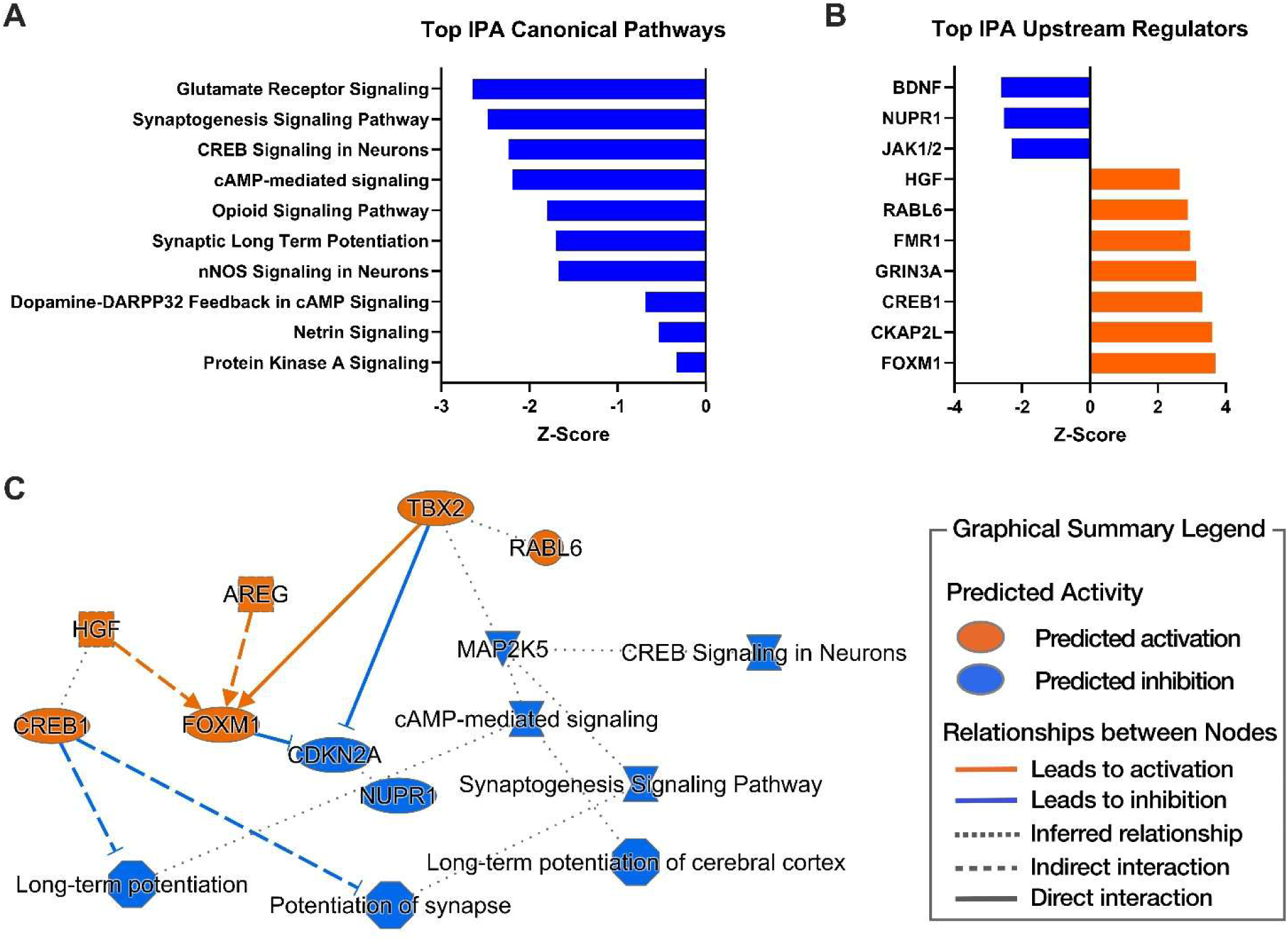
Dorsal Forebrain RNAseq analysis following treatment with 3 mg/kg. α**-cypermethrin from GD 6-16.** (A) Top 10 significantly altered canonical pathways in response to 3 mg/kg α-cypermethrin via Ingenuity Pathway analysis. (B) Top 10 predicted upstream regulators in response to 3 mg/kg cypermethrin via Ingenuity Pathway analysis. (C) Graphical summary of gene expression changes and node relationships produced by 3 mg/kg α-cypermethrin via Ingenuity Pathway Analysis. Bars in (A) and (B) represents predicted activation (orange) or inhibition (blue) of a pathway respectively. In (C), ovals signify transcriptional regulators, octagons signify biological functions, triangles signify kinases, and hour-glass shapes signify canonical pathways.

### Placental gene expression correlated with fetal outcomes

We also evaluated relationships between placenta gene expression and fetal brain development since our previous investigation demonstrated that pyrethroids are found in low levels in fetal tissue and may act through indirect, placental mechanisms (Elser et al. 2022). Using WGCNA to identify genes with co-regulated expression in placenta across vehicle and 10 mg/kg groups, we identified 17 clusters (i.e. modules) (Supplemental Figure 10B) and correlated the eigengenes of each module with cellular traits in matched fetal brains (Supplementary Figure 10A). Although no module-trait correlations survived correction after multiple testing, the lightgreen module positively correlated with percentage of ameboid microglia in the fetal dorsal forebrain, close to genome-wide significance (r = 0.63, FDR = 0.06, p = 0.004) (Supplemental Figure 10A). Hub genes of this module (Supplemental Figure 10C) that were linked through protein-protein networks (Supplemental Figure 10D) were also functionally enriched for biological processes such as cell chemotaxis (p= 7.13 x10^-7^), molecular functions (Supplemental Table 2) such as chemokine activity (p= 2.92 x10^-07^), and the chemokine signaling pathway (p = 3.17 x10^-07^) (Supplemental Table 3). This network was also enriched for DEGs (p < 0.004) which includes *Ccl3* and *Acod1* (aka *Irg1*). Together, this suggests that placental chemokine activity was linked with fetal brain microglia changes and that α-cypermethrin affected placental expression of genes present in this network.

### Doral forebrain RNAseq analysis

RNAseq analysis of the dorsal forebrain was performed to further assess cypermethrin-induced effects on neurodevelopment. Due to the potential confounding effect of maternal toxicity in response to 10 mg/kg α-cypermethrin and the lack of significant developmental alterations induced by permethrin, RNAseq analysis of the dorsal forebrain was performed to assess differences between the 3 mg/kg α-cypermethrin treated brains and controls. Bulk mRNA sequencing revealed 1,018 genes that were differentially expressed in response to 3 mg/kg α-cypermethrin (481 genes upregulated and 537 genes downregulated; Supplemental Figure 11). Pathway analysis implicated significant downregulation of c-AMP mediated signaling (z-score= −2.191, p=5.99E-11) and the synaptogenesis signaling pathway (z-score=-2.475, p=4.7E-07), among others (Figure 7). In addition, upstream regulator analysis in IPA predicted activation of CREB1 (z-score=3.3, p=1.54E-08), GRIN3A (z-score=3.1, p=2.41E-08), and CKAP2L signaling (z-score=3.6, p=1.16E07); as well as inhibition of BDNF signaling (z-score=-2.6, p=6.24E-08) (Figure 7).

### Weighted Gene Correlation network analysis (WGCNA) between placenta and fetal brain

To explore relationships between placenta and fetal dorsal forebrain gene expression, we performed WGCNA analysis using matched tissues exposed to vehicle or 3 mg/kg α-cypermethrin. We observed 31 modules of co-regulated genes from fetal brain (Figure 8A) and 22 modules from the placenta (Figure 8B). There was significant enrichment of fetal brain α-cypermethrin-induced DEGs in hub genes of multiple fetal brain modules (Table 3). In contrast, there was no significant enrichment of placenta α-cypermethrin-induced DEGs in hub genes of placenta modules.

**Fig. 8.**
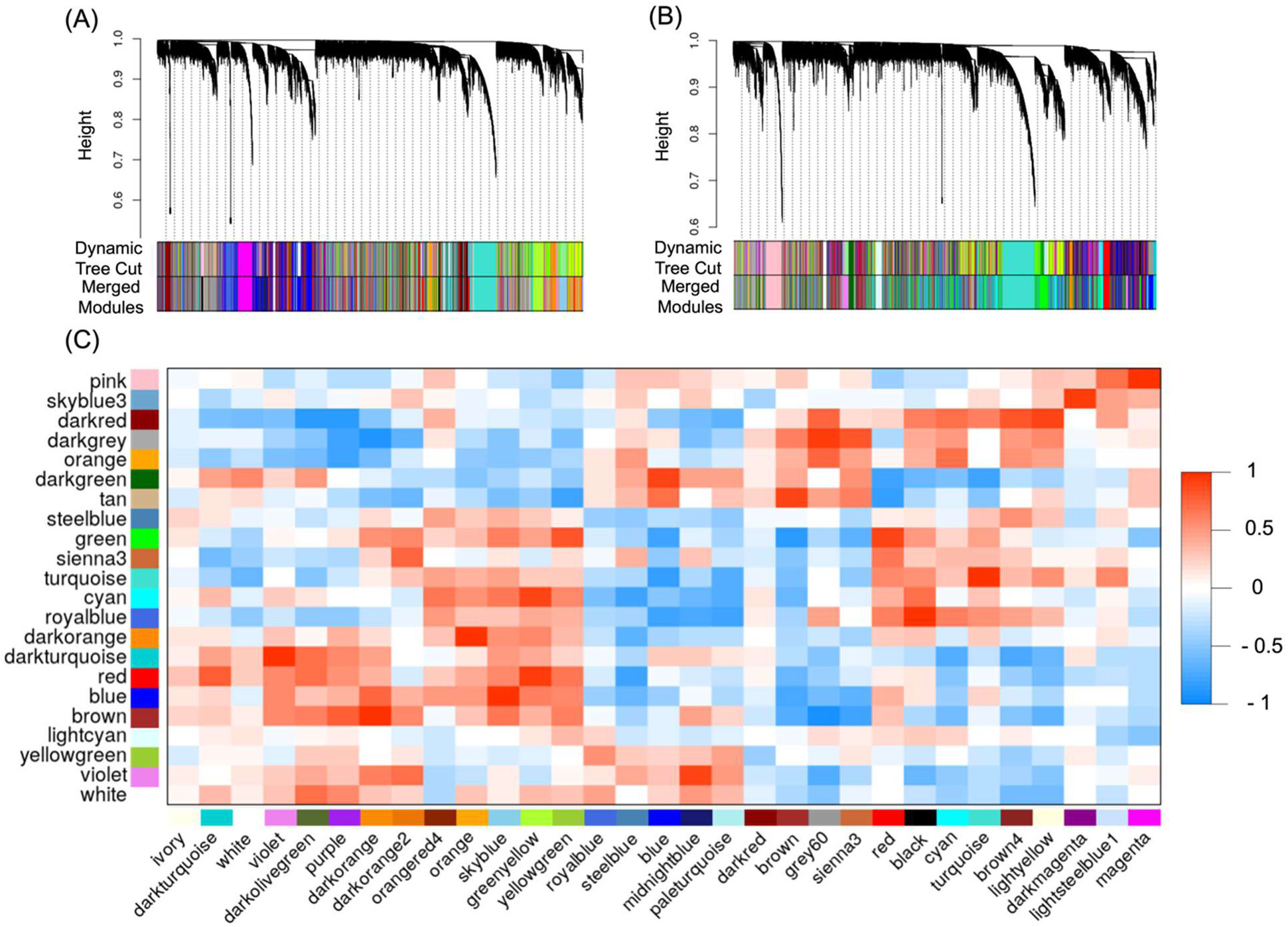
Correlation of eigengenes from fetal dorsal forebrain and placenta modules across vehicle and 3 mg/kg α-cypermethrin GD6-16 exposure. Hierarchical clustering was performed to identify modules from the (A) fetal dorsal forebrain and (B) placenta. (C) Pearson’s correlation was performed to evaluate the relationship between module eigengenes from the fetal brain (x-axis) and placenta (y-axis). Modules for each tissue were arbitrarily assigned a color name with some modules having the same name between the two tissues. There is however no intention of suggesting any relationship between these modules with the same name. Increasing red indicates increasing positive correlation while increasing blue indicates increasing negative correlation.

Using a representative value of each module (eigengene values) of each tissue, we identified module pairs that had correlated expression between the different tissues (Figure 8C). To examine these relationships, we further analyzed the three fetal brain modules with greatest enrichment for α-cypermethrin-induced DEGs-darkorange (odds ratio = 12.9; FDR = 6.2 x10^-129^), grey60 (odds ratio = 10.9; FDR = 1.4 x10^-111^) and midnightblue (odds ratio = 11.4; FDR = 6.61 x10^-77^) (Table 3). Gene ontology analysis using hub genes of these modules showed significant enrichment for biological processes involved in regulation of neurotransmitter regulation, transport and secretion for the darkorange module (Figure 9A top panel, Table 4), regulation of mitosis including mitotic nuclear division and regulation of mitotic cell cycle for the grey60 module (Figure 9B top panel, Table 4), and synapse organization for the midnightblue module (Figure 9C top panel, Table 4).

**Fig. 9.**
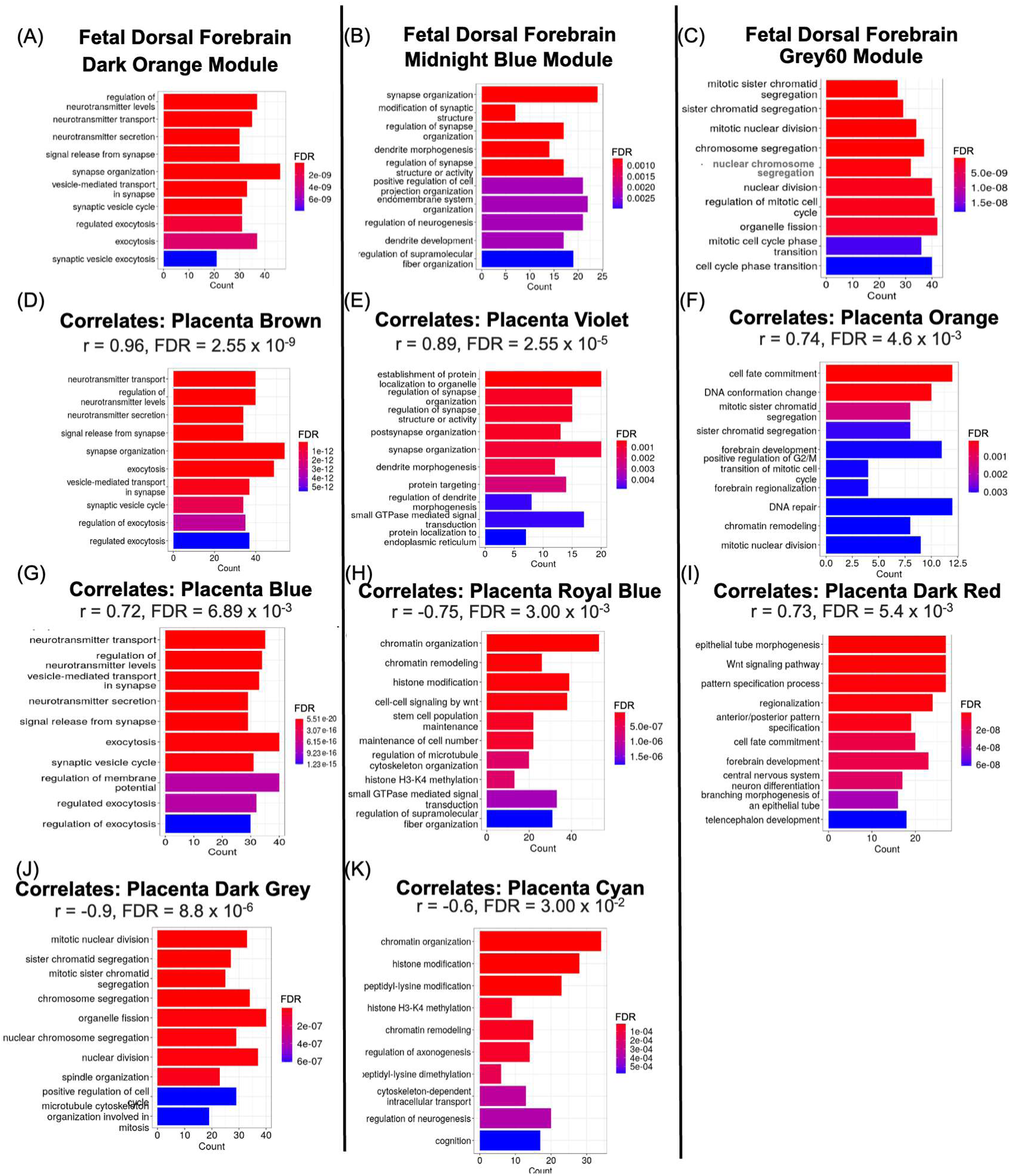
Gene ontology analysis of biological processes for fetal dorsal forebrain and corresponding placenta modules that correlates with it. Fetal brain modules with greatest enrichment of DEGs from α-cypermethrin exposure were enriched for particular biological processes (A-C) that were similar to those enriched in the placenta modules (D-K) that significantly correlated with these fetal brain modules through gene ontology analysis. Each panel describes the tissue and modules associated with that tissue. For placenta modules (D-K), the panel title includes the correlation value and significance of that correlation with the fetal brain module directly above. The top 10 biological processes are shown. Please refer to Table 6 for a full list of correlations between fetal brain and placenta. r; Pearson’s correlation coefficient between fetal brain and placenta modules, FDR; false discovery rate used to correct for multiple testing.

Interestingly, these three fetal dorsal forebrain modules were highly correlated with several placenta modules (Table 5). Notably, the fetal brain darkorange module showed significant correlation with the placenta brown (r = 0.96, FDR = 2.55 x 10^-9^) and blue (r = 0.72, FDR = 6.89 x 10^-3^) modules, which were similarly significantly enriched for biological processes including neurotransmitter regulation, secretion, and transport (Figure 9D,G, Table 6). In contrast, it was negatively correlated with the darkgrey placenta module (r = −0.9, FDR = 8.8 x 10^-6^) which was enriched for biological processes involved in mitosis and cell division (Figure 9J, Table 6). The fetal brain midnightblue module positively correlated with the placenta violet module (r = 0.89, FDR = = 1.21 x 10^-5^), also enriched for biological processes involved in synapse organization and activity (Figure 9E). The fetal brain midnight blue module was negatively correlated with placental royal blue (r = −0.75, FDR = 0.003) and cyan (r = −0.60, FDR = 0.047) modules (Figure 9E,H,C, Table 6) and enriched for biological processes involved in chromatin remodeling (Figure 9E,H). The fetal brain grey60 module correlated with the orange (r = 0.74, FDR = 0.0046) and darkred (r = 0.73, FDR = 0.0054) placenta modules which were enriched for processes including those involved in, respectively, cell fate development and mitosis and Wnt signaling and forebrain and central nervous system development (Figure 9F,I). Gene ontology analysis of other modules are detailed in Table 6.

Given that modules of correlated gene expression across tissues shared similar biological processes from gene ontology analysis, we further investigated if there was a relationship between these two factors: the degree of overlap in biological processes and the strength of correlation between brain and placenta modules that correlated. Using all module-pairs that were significantly correlated and had biological process enrichment, we overall observed a moderate but significant correlation between the two factors (r = 0.44, p = 0.00012) (Figure 10). This suggests that the correlation strength between modules of each tissue could be due to the overlapping biological processes shared between them.

**Fig. 10.**
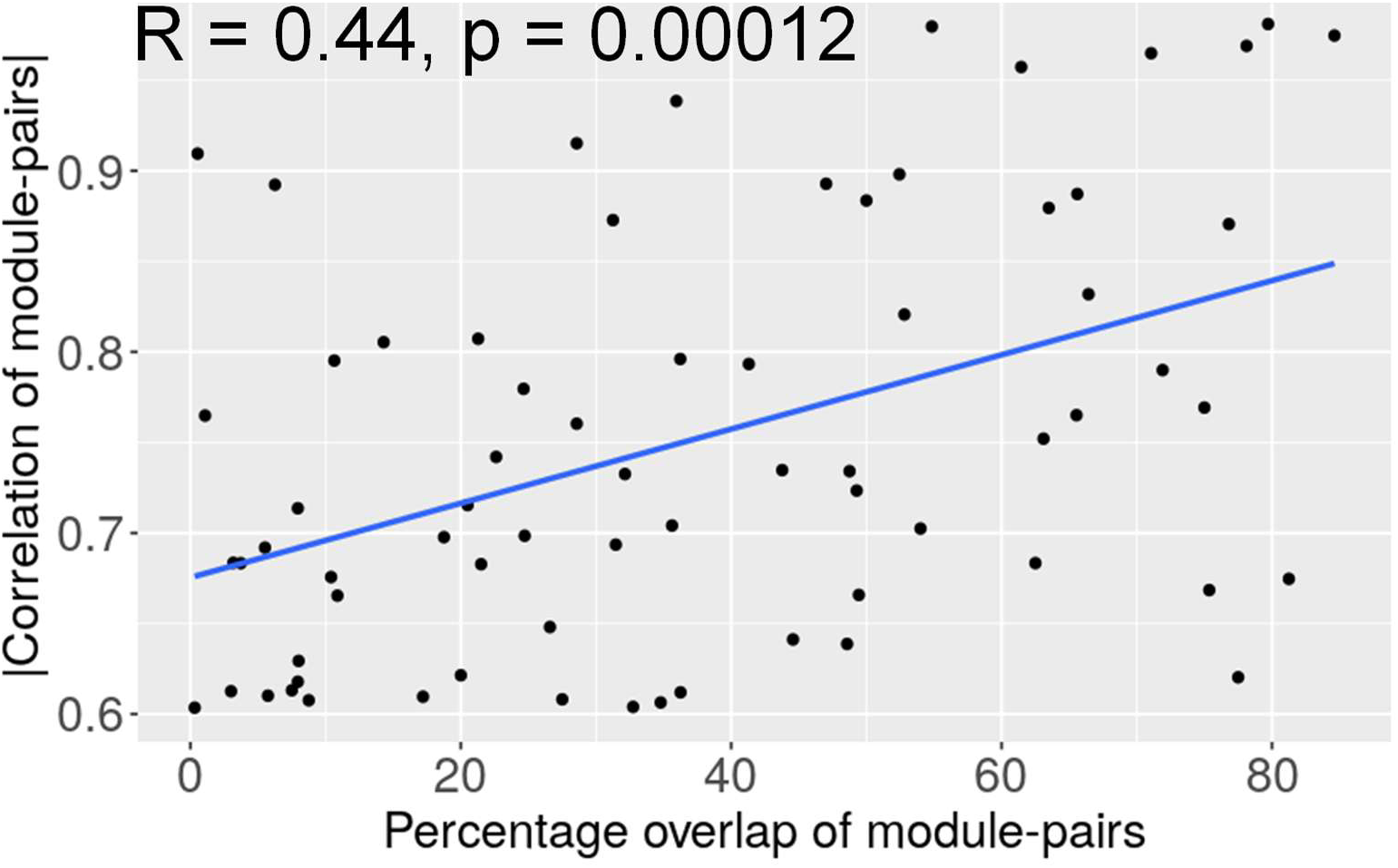
Relationship between fetal brain to placenta module extent of correlation and percentage of overlapping biological processes of module-pairs. Pearson correlation was performed to evaluate the correlation of module-pairs across brain and placenta coregulated gene modules (|correlation of module-pairs|) and the percentage of overlapping biological processes that these genes were enriched for. Data consists of seventy module-pairs that were significantly correlated between fetal brain and placenta.

## Discussion

Overall, results presented from this study implicate the placenta as a potentially relevant target organ for type II pyrethroid exposure in pregnancy. We observed a dose-dependent decrease in mouse fetal body weight specifically for the type II pyrethroid α-cypermethrin, despite no effects on fetal growth for the type I pyrethroid permethrin. In addition, we report alterations in placental layer morphology in response to α-cypermethrin exposure that are consistent with models of growth restriction, despite no effects on placental morphology for permethrin. Given the absence of effects on maternal body weight, liver pathology, or inflammatory markers, our data suggest that a maternally-mediated mode of action is a less-likely contributor to the effects of α-cypermethrin on fetal growth within this study.

The placental labyrinth zone is comprised of maternal and fetal blood vessels that serve as the site of nutrient and oxygen exchange between mother and fetus. Impairments in the development of this zone can cause intrauterine growth restriction through reduced blood flow and nutrient supply to the fetus (Perez-Garcia et al. 2018; Woods, Perez-Garcia, and Hemberger 2018). In our study, we observed a dose-dependent reduction the size of the labyrinth zone in response to α-cypermethrin, which correlated with effects on fetal growth. Pathway analysis of placental RNAseq data revealed downregulation of genes involved in extracellular matrix (ECM) organization in response to α-cypermethrin, a process that facilitates the morphogenesis of placental tissue to support a healthy pregnancy. Dysregulation of ECM remodeling is associated with pregnancy-specific disease states that reduce fetal growth, such as preeclampsia (O’Connor et al. 2020), and can impact critical processes such as cell migration and angiogenesis (Bonnans, Chou, and Werb 2014; Sottile 2004). As such, it is possible that effects reported on labyrinth zone development could be the result of alterations in extracellular matrix remodeling. With respect to the lack of effects reported for permethrin, it is also worth considering the role that chloride channels play in regulating angiogenesis (Li et al. 2021; Kamili et al. 2020). Cypermethrin and other type II pyrethroids inhibit chloride channel opening, while permethrin and most other type I pyrethroids have no significant effect on these channels (Breckenridge et al. 2009). Previous reports have demonstrated that chloride channel inhibition impairs angiogenesis, and as such, the difference in effects could be explained through this mechanism (Kamili et al. 2020).

We also report effects of α-cypermethrin on the placental junctional zone, the endocrine compartment of the placenta responsible for producing growth factors and hormones critical for fetal development. The junctional zone comprises three distinct cell types, including spongiotrophoblasts, glycogen cells, and trophoblast giant cells. Spongiotrophoblasts are implicated mainly in the production of hormones, while glycogen cells are suspected of playing a role in the supply of glucose to the fetus (Woods, Perez-Garcia, and Hemberger 2018). Our study found that α-cypermethrin exposure resulted in a larger junctional zone with an increase in glycogen cells, a phenotype implicated in several models of growth restriction (Tunster et al. 2020; Tunster et al. 2016; Tunster, Van de Pette, and John 2012; Esquiliano et al. 2009). Increased numbers of glycogen cells within the junctional zone may reflect abnormalities in several processes, including altered differentiation of these cells, impairments in the degradation of glycogen, or mis-localization of these cells resulting from impairments to their migration (Woods, Perez-Garcia, and Hemberger 2018).

Abnormal junctional zone development observed here may also be explained by alterations in genes involved in ECM remodeling, as this process facilitates the migration of placental glycogen cells from the junctional zone to the decidua (Jia et al. 2014; Yu et al. 2014; Ding, Huang, et al. 2015; Coan et al. 2006). Further investigation into ECM-related changes revealed a significant downregulation of placental *Lox* expression in response to α-cypermethrin and an increase in placental collagen staining within the junctional zone. These results are consistent with previous work demonstrating that downregulation of placental *Lox* expression impairs trophoblast migration and increases trophoblast collagen production through activation of the TGF-β1/Smad3/collagen pathway, as seen in cases of preeclampsia (Xu et al. 2019). While the consequences of impaired migration of glycogen cells are unclear, their association with maternal blood spaces within the decidua suggests that altered migration may impact glucose release into fetal circulation (Woods, Perez-Garcia, and Hemberger 2018).

In addition to effects on placenta, our data also demonstrate alterations in fetal dorsal forebrain development with maternal pyrethroid exposure. Results from this study build upon our previous findings (Elser et al. 2020) by demonstrating a dose-dependent shift in fetal microglia morphology from a mature ramified state to an immature ameboid state following maternal α-cypermethrin exposure throughout organogenesis (GD 6 to 16). These dose-dependent effects were apparent even at the lowest dose of α-cypermethrin (0.3 mg/kg), which produced blood levels of α-cypermethrin in pregnant mice within the range of environmental exposures in humans (Elser et al. 2022). Permethrin showed similar dose-dependent alterations in microglia morphology, but at a lower magnitude than those induced by α-cypermethrin. In general, our data suggest that the shift in microglial morphology may result from a developmental delay. However, the increase in multivacuolated morphology observed at ≥ 3 mg/kg α-cypermethrin may also indicate an activated state of these cells.

Pathway analysis of fetal brain gene expression changes in response to maternal exposure to 3 mg/kg α-cypermethrin implicated alterations in genes downstream of BDNF signaling, as well as alterations in neurodevelopmental processes such as synaptogenesis. Notably, microglia promote synaptogenesis through BDNF signaling (Parkhurst et al. 2013), so the alterations in microglia development may be linked to changes in synaptic development suggested by our RNAseq data. Pathway analysis also suggested a cypermethrin-induced downregulation of c-AMP signaling, which has been implicated in the regulation of microglial process extension in the adult brain (Bernier et al. 2019; Ghosh, Xu, and Pearse 2016).

Given that we found relatively low levels of each pyrethroid within the fetus (Elser et al. 2022), we also investigated whether effects on microglial development could be tied to alterations in placental function. Interestingly, we observed that a placental gene network tied to chemokine activity was associated with ameboid microglia density, suggesting a possible role of the placenta in mediating effects of α-cypermethrin on microglial development. While it is presently unclear how placental chemokine activity affects fetal brain microglia morphology, chemokines such as Ccl5 and Ccl3 regulate processes such as trophoblast invasiveness and angiogenesis within the placenta, which impact fetal brain development (Du, Wang, and Li 2014).

Through correlational analysis of co-regulated genes in placenta and brain, we also identified biological processes in the placenta that had strong relationships with processes in the fetal brain. For example, expression levels of co-regulated gene modules involved in neurotransmitter regulation and synapse organization were affected by α-cypermethrin and positively correlated between placenta and fetal brain. This is consistent with previous work demonstrating that neurotransmitters secreted from the placenta regulate fetal brain cell division, neuronal migration, differentiation, and synaptogenesis (Rosenfeld 2021). Similarly, alterations to *Wnt* signaling can affect development of both the placenta and fetal brain (Dietrich et al. 2022; Knöfler and Pollheimer 2013; Akram et al. 2022). As such, processes which regulate placenta development and function may also impact fetal brain development across a range of biological domains.

In addition to these findings, our RNAseq workflow identified several covariates that may be useful for incorporation into future studies assessing fetal transcriptomics. In placenta and dorsal forebrain RNAseq results, we identified several covariates that significantly affected gene expression, including the breeder male, litter size, sex, and uterine position relative to the cervix. It is interesting to note the significant effect that the four sibling breeder males used in this study had on placental and fetal brain gene expression, as their similar age and genetic background suggest potential epigenetic paternal contributions to fetal gene expression. Effects that uterine position had on fetal gene expression was particularly interesting in the context of previous work suggesting that uterine position may affect placental blood flow and fetal exposure to toxicants (Even et al. 1994; McLaurin and Mactutus 2015). Overall, incorporating these factors into our analysis model enhanced our ability to detect transcriptional changes induced by pyrethroid exposure. Based on our findings, we recommend that future studies investigating fetal gene expression changes consider these covariates to avoid type I and II errors due to their effects.

In summary, our study demonstrated dose-dependent alterations in fetal growth, placental development, and neurodevelopment in response to the type II pyrethroid α-cypermethrin, despite minimal effects on these endpoints by the type I pyrethroid permethrin. These data suggest that the placenta may be a relevant target organ of type II pyrethroids and that further investigations should be performed to elucidate mechanisms by which alterations in placental development may occur. In addition, our study implicates dose-dependent microglia developmental delays, which may affect many neurodevelopmental processes. Overall, these data suggest increased concern specifically for type II pyrethroids in regard to embryo-fetal development. As such, more work should be done to clarify whether this distinction between type I and II pyrethroids is relevant to reported effects on fetal development in humans.

## Supporting information

All Supplemental Figures and Tables

## Acknowledgements

RNA sequencing was performed with the support of the Iowa Institute of Human Genetics Core. Placenta RNAseq reads were mapped to a reference genome with the support of the University of Iowa Bioinformatics Division.

## Supplementary Materials

The supplementary materials consist of 11 figures and 3 tables. Figure 1: Liver histology was unaffected by 10 mg/kg α-cypermethrin or 50 mg/kg permethrin. Figure 2: Cypermethrin alters placental layer thickness. Figure 3: Correlational analyses between fetal weight, dorsal forebrain volume, and labyrinth zone thickness. Figure 4: Volcano plots of differential gene expression in the placenta. Figure 5: Top gene expression pathways in GD 16 mouse placenta altered by 50 mg/kg permethrin (iPathway). Figure 6: Evaluation of oxidative stress in pyrethroid-treated placentas. Figure 7: Effects of pyrethroids on mouse placental macrophage density and cytokine production. Figure 8: Sex-differences in the effects of cypermethrin and permethrin on ramified microglia in the GD 16 fetal mouse brain. Figure 9: P2ry12 staining and gene expression of P2ry12, Cd68, Apoe, and Il-1b in GD 16 mouse fetal brain. Figure 10: Weighted gene correlation network analysis (WGCNA) of placental RNAseq. Figure 11: Volcano plot of differential gene expression in the dorsal forebrain. Table 1: Primer Sequences used in qPCR assays. Table 2: Placental RNAseq WGCNA molecular functions. Table 3: Placental RNAseq WGCNA pathway analysis.

