## Supplementary material for "Maternal α-cypermethrin and permethrin exert differential effects on fetal growth, placental morphology, and fetal neurodevelopment in mice": All Supplemental Figures and Tables

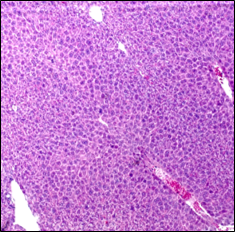

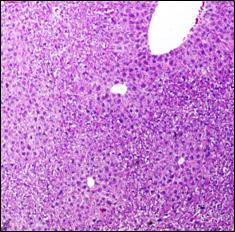

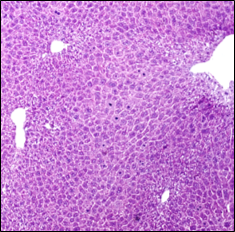

**Vehicle**

**Cyp 10 mg/kg**

**Per 50 mg/kg**

Supplemental Figure 1. Liver histology was unaffected by 10 mg/kg α-cypermethrin or 50 mg/kg permethrin. Representative pictures of livers showing no signs of overt toxicity. n=5 livers per treatment group were evaluated by a board-certified pathologist.

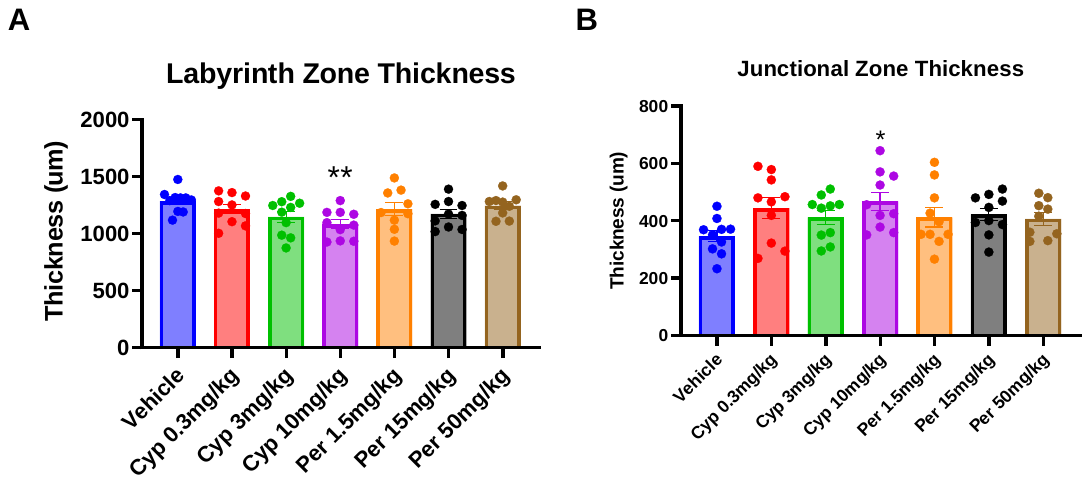

Supplemental Figure 2. Cypermethrin alters placental layer thickness. (A) Cypermethrin decreased the thickness of the placental labyrinth zone. (B) Cypermethrin increased the thickness of the placental junctional zone. Means ± SEMs are shown. n=9-10 litters per treatment group. (*p<0.05, **p<0.01 by post hoc tests) Cyp= Cypermethrin, Per= Permethrin.

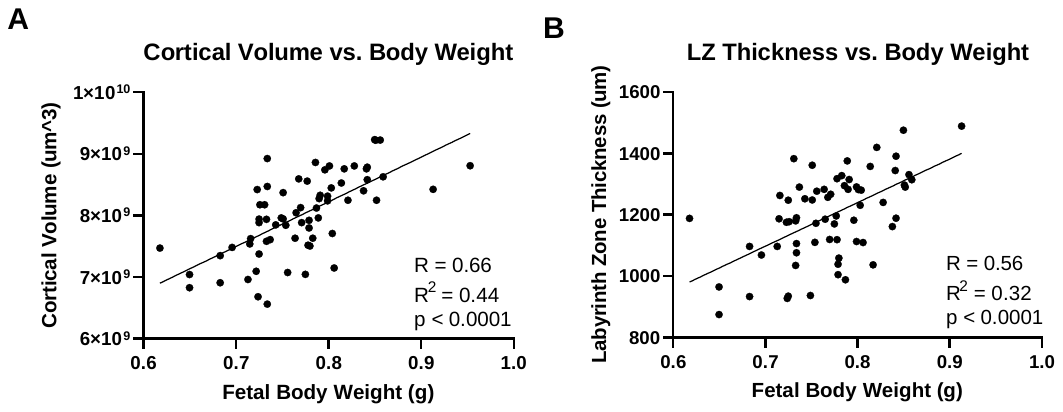

**Dorsal Forebrain Volume vs Body Weight**

Supplemental Figure 3. Correlational analyses between fetal weight, dorsal forebrain volume, and labyrinth zone thickness. (A). Fetal body weight correlates with dorsal forebrain volume. (B) Fetal body weight correlates with placental labyrinth zone thickness.

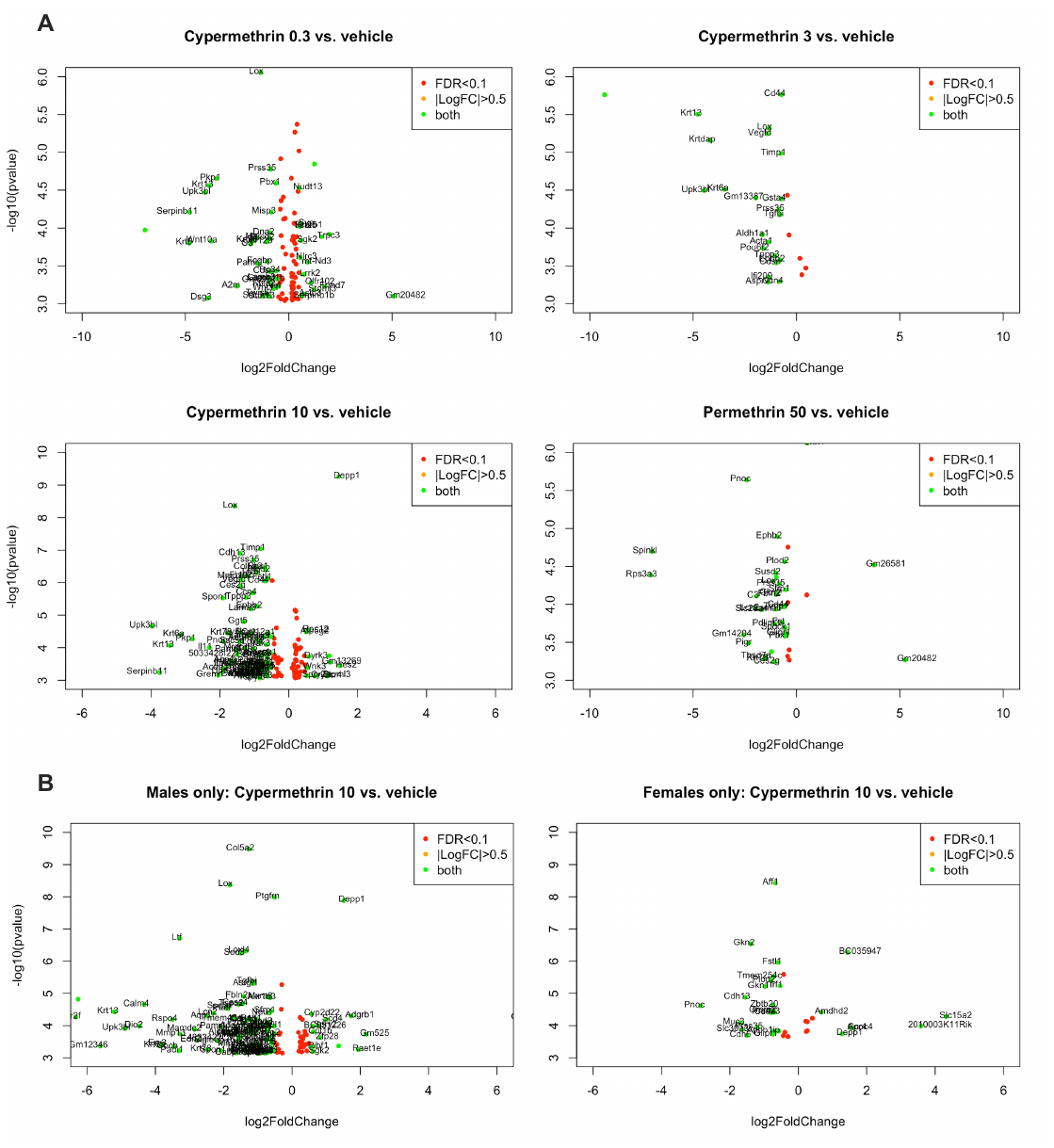

Supplemental Figure 4. Volcano plots of differential gene expression in the placenta.

Positive log2 fold change indicates higher gene expression compared to controls. Negative log2 fold change indicates lower gene expression compared to controls. Green indicates genes with log2 fold change > 0.5 and FDR < 0.1.

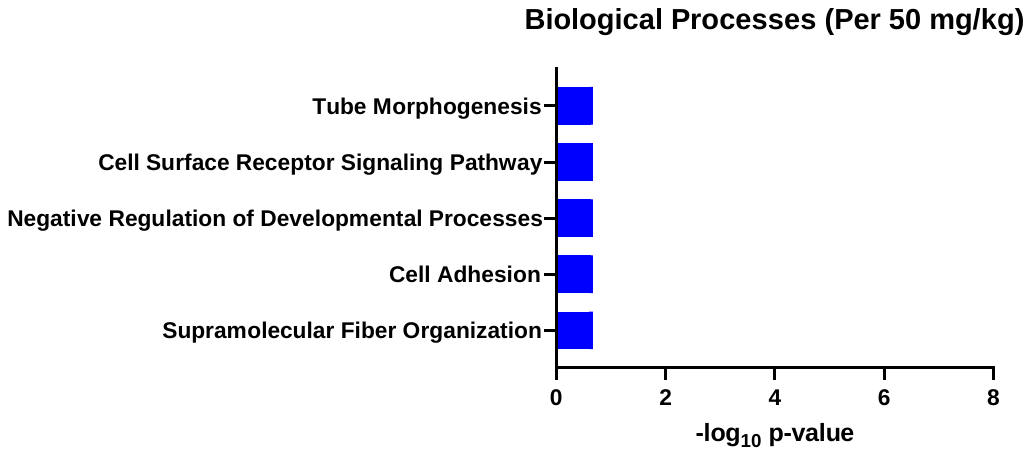

Supplemental Figure 5. Top gene expression pathways in GD 16 mouse placenta altered by 50 mg/kg permethrin (iPathway).

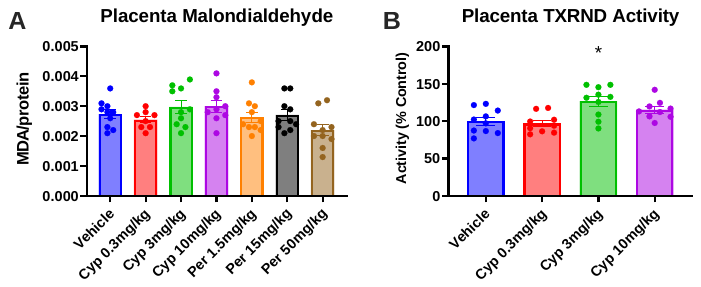

Supplemental Figure 6. Evaluation of oxidative stress in pyrethroid-treated placentas.

(A) Neither pyrethroid significantly altered placental malondialdehyde content. (B) 3 mg/kg cypermethrin increased TNXRD activity in response to cypermethrin exposure. Means ± SEMs are shown. n=9-10 litters per treatment group. (*p<0.05). Cyp= Cypermethrin, Per= Permethrin.

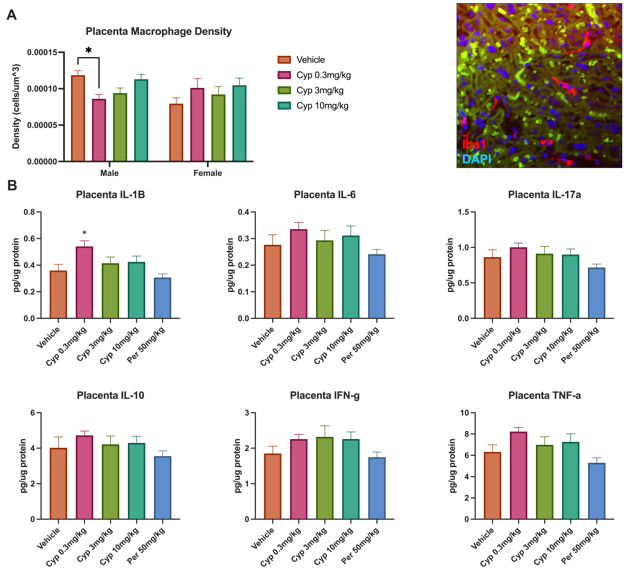

Supplemental Figure 7. Effects of pyrethroids on mouse placental macrophage density and cytokine production.

(A) 0.3 mg/kg cypermethrin decreased placental macrophage density in males specifically. Representative picture of placental Iba1 staining on right. (B) 0.3mg/kg cypermethrin increased placental IL-1B, however, no other effects were reported for either pyrethroid on placental cytokines. Means ± SEMs are shown. n=8-10 litters per treatment group. *p<0.05 by post hoc test). Cyp= Cypermethrin, Per= Permethrin.

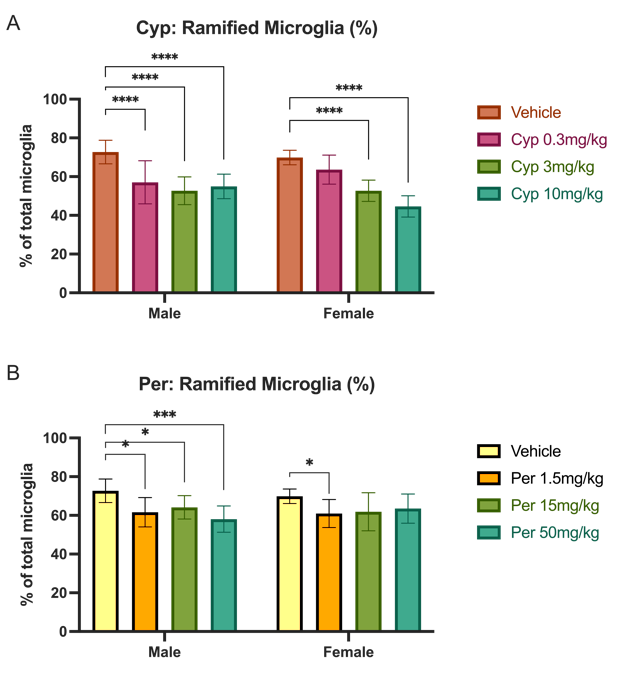

Supplemental Figure 8. Sex-differences in the effects of cypermethrin and permethrin on ramified microglia in the GD 16 fetal mouse brain.

(A) A significant interaction between cypermethrin and sex was observed with regard to the percentage of ramified microglia (p=0.0049). Data are shown separated by sex. (B) No significant interaction was observed between permethrin and sex with regard to percentage of ramified microglia. Means ± SDs are shown. n=6-10 brains/sex/treatment. (*p<0.05, ***p<0.001, ****p<0.0001 by post hoc tests). Cyp= Cypermethrin, Per= Permethrin.

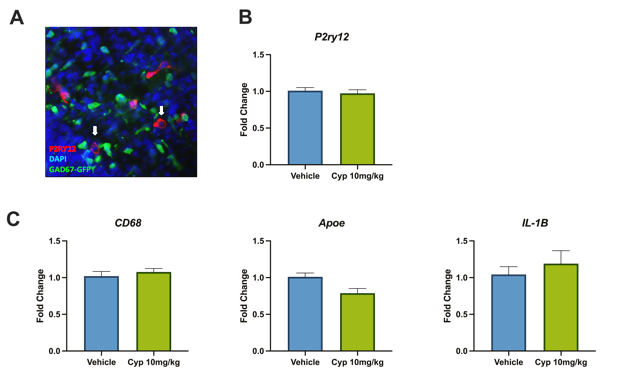

Supplemental Figure 9. P2ry12 staining and gene expression of P2ry12, Cd68, Apoe, and Il-1b in GD 16 mouse fetal brain. (A) Representative image of P2ry12 staining demonstrating labeling of ameboid morphology. (B) Cypermethrin did not alter gene expression of microglia resting state marker P2ry12in the fetal brain. (C) Cypermethrin did not significantly alter gene expression of microglia activation state markers in the fetal brain. Means ± SEMs are shown. n=9-10 litters per treatment group. Cyp= Cypermethrin.

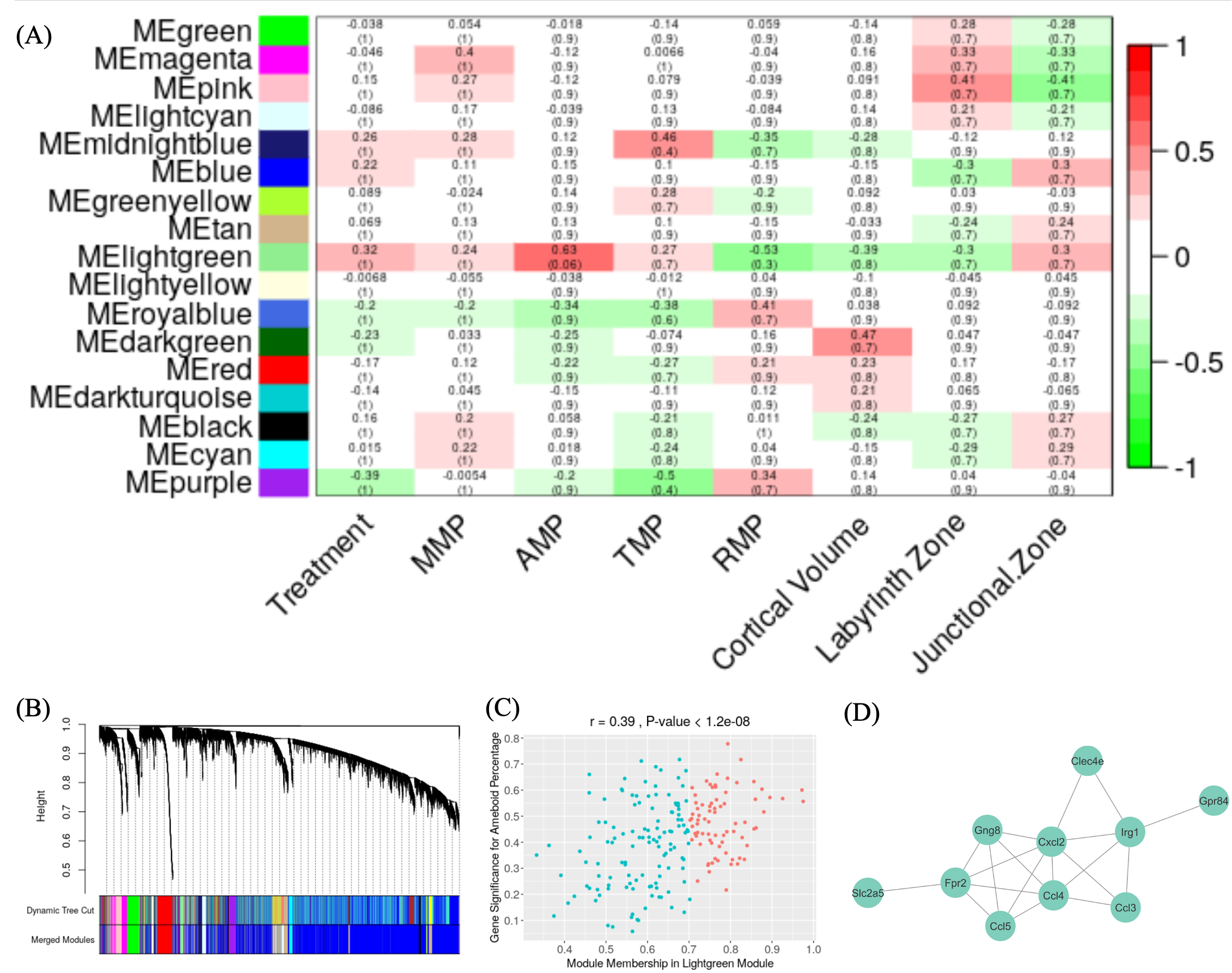

Supplemental Figure 10. Weighted gene correlation network analysis (WGCNA) of placental RNAseq.

(A) Module-trait correlation analysis. Modules from placental RNAseq were identified using WGCNA.

Correlation was performed between eigengenes from each module and different conditions such as treatment (control or cypermethrin), multivacuolated microglia percentage (MMP), ameboid microglia percentage (AMP), transitional microglia percentage (TMP), ramified microglia percentage (RMP), cortical volume, area of placenta labyrinth zone (Labyrinth Zone) and area of placenta junctional zone (Junctional Zone). Color scale represent module-trait correlation from -1 (green) to 1 (red). Number in the table reflects Pearson’s correlation coefficient and numbers in braces reflects FDR corrected P-values. (B) Dendrogram of genes clustered together based on similarity of their expression profile forming gene modules that were assigned different colors. Modules with genes that had highly similar co-expression were merged into a single module (Merged Modules). (C) Identification of hub genes (colored in red) defined by Gene Significance ≥ 0.2 and Module Membership ≥ 0.7. Genes which are not hub genes are colored in cyan. (D) Protein-protein network identified from hub genes of the lightgreen module.

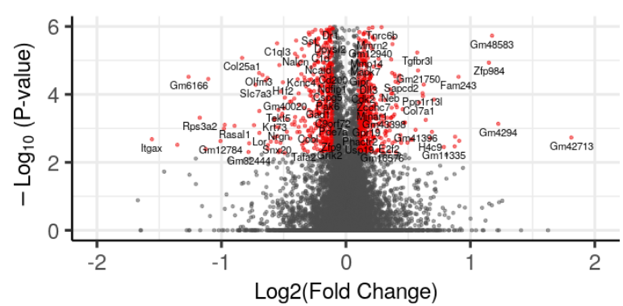

Supplemental Figure 11. Volcano plot of differential gene expression in the dorsal forebrain.

Positive log2 fold change indicates higher gene expression from embryonic forebrain with prenatal cypermethrin exposure compared to controls. Negative log2 fold change indicates lower gene expression from embryonic forebrain with prenatal exposure to cypermethrin compared to controls. Red indicates genes with log2 fold change > 0.1 and FDR < 0.05.

Supplemental Table 1. Primer sequences used in qPCR assays

| **Supplementary Table 1. Primer sequences used in qPCR assays** | | |
| --- | --- | --- |
| **Gene** | **Forward Primer** | **Reverse Primer** |
| **Apoe** | ATTGCTGACAGGATGCCTAGC | GGTTGGTTGCTTTGCCACTC |
| **Cd68** | TCCAAGATCCTCCACTGTTG | ATTTGAATTTGGGCTTGGAG |
| **Depp1** | TGAGCACTCTCTGGGAAGAAAAC | GATCACTGGGAGGTGCAAATAGA |
| **Gapdh** | GGTGAAGGTCGGTGTGAACG | CTCGCTCCTGGAAGATGGTG |
| **Il-1b** | TTCACCATGGAATCCGTGTC | GTCTTGGCCGAGGACTAAGG |
| **Lox** | CAGCCACATAGATCGCATGGT | GCCGTATCCAGGTCGGTTC |
| **P2ry12** | TTCCTGGGGTTGATAACCATTG | GGTGAGAATCATGTTAGGCAGTG |
| **Sdha** | GCTCCTGCCTCTGTGGTTGA | AGCAACACCGATGAGCCTG |

Supplemental Table 2. Placental RNAseq WGCNA molecular functions

| #term ID | term description | observed gene count | background gene count | strength | FDR |
| --- | --- | --- | --- | --- | --- |
| GO:0008009 | chemokine activity | 4 | 41 | 2.33 | 2.92E-07 |
| GO:0048020 | CCR chemokine receptor binding | 3 | 35 | 2.28 | 1.74E-05 |
| GO:0031726 | CCR1 chemokine receptor binding | 2 | 4 | 3.04 | 5.16E-05 |
| GO:0031730 | CCR5 chemokine receptor binding | 2 | 7 | 2.8 | 7.73E-05 |
| GO:0016004 | phospholipase activator activity | 2 | 11 | 2.6 | 0.00015 |
| GO:0042056 | chemoattractant activity | 2 | 29 | 2.18 | 0.00061 |
| GO:0005102 | signaling receptor binding | 5 | 1515 | 0.86 | 0.0019 |
| GO:0008528 | G protein-coupled peptide receptor activity | 2 | 128 | 1.54 | 0.0093 |
| GO:0030246 | carbohydrate binding | 2 | 265 | 1.22 | 0.0318 |
| GO:0038023 | signaling receptor activity | 3 | 1020 | 0.81 | 0.0457 |

Supplemental Table 3. Placental RNAseq WGCNA pathway analysis

| #term ID | term description | observed gene count | background gene count | strength | FDR |
| --- | --- | --- | --- | --- | --- |
| mmu04062 | Chemokine signaling pathway | 5 | 179 | 1.79 | 3.17E-07 |
| mmu04060 | Cytokine-cytokine receptor interaction | 4 | 252 | 1.54 | 5.99E-05 |
| mmu05132 | Salmonella infection | 3 | 78 | 1.93 | 6.37E-05 |
| mmu04620 | Toll-like receptor signaling pathway | 3 | 98 | 1.83 | 9.28E-05 |
| mmu04623 | Cytosolic DNA-sensing pathway | 2 | 61 | 1.86 | 0.0024 |
| mmu05323 | Rheumatoid arthritis | 2 | 81 | 1.74 | 0.0035 |
| mmu04064 | NF-kappa B signaling pathway | 2 | 93 | 1.68 | 0.0039 |
| mmu05142 | Chagas disease (American trypanosomiasis) | 2 | 101 | 1.64 | 0.004 |
| mmu04668 | TNF signaling pathway | 2 | 108 | 1.61 | 0.0041 |
| mmu04621 | NOD-like receptor signaling pathway | 2 | 164 | 1.43 | 0.0083 |
| mmu05167 | Kaposi's sarcoma-associated herpesvirus infection | 2 | 203 | 1.34 | 0.0114 |
